# Prophase removal of chromosome-associated RNAs facilitates anaphase chromosome segregation

**DOI:** 10.1101/813527

**Authors:** Judith A. Sharp, Wei Wang, Michael D. Blower

## Abstract

During mitosis, the genome is transformed from a decondensed, transcriptionally active state to a highly condensed, transcriptionally inactive state. Mitotic chromosome reorganization is marked by the general attenuation of transcription on chromosome arms, yet how the cell regulates nuclear and chromatin-associated RNAs after chromosome condensation and nuclear envelope breakdown is unknown. SAF-A/hnRNPU is an abundant nuclear protein with RNA-to-DNA tethering activity, coordinated by two spatially distinct nucleic acid binding domains. Here we show that RNA is evicted from prophase chromosomes through Aurora-B-dependent phosphorylation of the SAF-A DNA-binding domain; failure to execute this pathway leads to accumulation of SAF-A:RNA complexes on mitotic chromosomes and elevated rates of anaphase segregation defects. This work reveals a role for Aurora-B in removing chromatin-associated RNAs during prophase, and demonstrates that Aurora-B dependent relocalization of SAF-A during cell division contributes to the fidelity of chromosome segregation.

## Introduction

During cell division, chromosomes are dramatically restructured to facilitate segregation to daughter cells. At the onset of mitosis, most nuclear transcription ceases through a block to transcription initiation, causing a run-off of actively elongating RNA polymerase (Akoulitchev & Reinberg, 1998; Liang et al., 2015; Palozola et al., 2017; Prescott & Bender, 1962; Segil, Guermah, Hoffmann, Roeder, & Heintz, 1996). During the same period, interphase chromosome architecture is erased through the removal of the majority of cohesin complexes from euchromatin and the loading of condensin complexes (Gibcus et al., 2018). This coordinated exchange of cohesin and condensin complexes leads to dramatic chromosome condensation and the characteristic X shape of mitotic chromosomes (Haarhuis, Elbatsh, & Rowland, 2014). Collectively, each of these genome remodeling pathways contributes to the accurate segregation of chromosomes during anaphase.

Aurora-B is a master regulatory kinase that controls mitotic chromosome segregation and cytokinesis. Aurora-B is a member of the ‘Chromosome Passenger Complex (CPC)’ that dynamically changes localization throughout mitosis. The CPC localizes throughout the chromosomes at prophase, concentrates at the inner centromere region at metaphase, and transfers from chromosomes to the spindle midzone during anaphase (Carmena, Wheelock, Funabiki, & Earnshaw, 2012). During prophase the CPC, Plk1, and Cdk1 phosphorylate the cohesin-associated proteins Sororin and SA2, which allows the cohesin release factor WAPL to open cohesin rings and strip cohesin from euchromatic regions (Haarhuis et al., 2014). In addition to phosphorylation of cohesin complexes, Aurora-B phosphorylates histone H3 during prophase to release HP1 from chromatin (Fischle et al., 2005; Hirota, Lipp, Toh, & Peters, 2005). However, the full spectrum of mitotic Aurora-B substrates and functions is not currently known.

During interphase a large proportion of the genome is transcribed into RNA. Most mRNAs are spliced, capped, polyadenylated and exported from the nucleus. However, a small fraction of fully processed mRNAs are retained in the nucleus (Bahar Halpern et al., 2015). The vast majority of nuclear transcripts do not code for proteins but are part of a diverse group of functional noncoding RNAs (Djebali et al., 2012). Noncoding RNAs influence gene expression by many mechanisms, including interacting with both transcriptional activators and repressors, promoting 3D genome organization, coating specific chromosomal domains, promoting DNA replication origin usage, and silencing an entire chromosome (Wang & Chang, 2011). Since noncoding RNA function has been studied primarily during interphase, it is not clear how genome organization promoted by nuclear RNAs is regulated during mitosis when chromosomes are restructured.

The prototypical nuclear noncoding RNA is the XIST RNA. This transcript is expressed from the inactive X chromosome in female cells and coats the Xi chromosome in *cis* to silence most genes (Galupa & Heard, 2015). XIST RNA is tethered to the Xi chromosome during interphase through a combination of factors including Ciz1 (Ridings-Figueroa et al., 2017; Sunwoo, Colognori, Froberg, Jeon, & Lee, 2017), hnRNP-K (Colognori, Sunwoo, Kriz, Wang, & Lee, 2019; Pintacuda et al., 2017), and hnRNP-U/SAF-A (Hasegawa et al., 2010). Interestingly, XIST RNA is removed from the mitotic Xi chromosome in an Aurora-B-dependent manner, but the molecular mechanism is not known (Hall, Byron, Pageau, & Lawrence, 2009). Additionally, it is not known if other nuclear RNAs are also removed from chromosomes during mitosis.

SAF-A/hnRNPU is a highly abundant nuclear protein originally identified as a factor that bound with high affinity to Scaffold Attachment Regions of chromosomes and as a protein that bound to hnRNA (Fackelmayer & Richter, 1994; Kiledjian & Dreyfuss, 1992). Indeed, SAF-A has been shown to interact with hundreds of RNAs, and recent mass spectrometry studies have shown that all cellular SAF-A is complexed with RNA (Caudron-Herger et al., 2019; Huelga et al., 2012; R. Xiao et al., 2012). SAF-A contains a modular domain architecture with a N-terminal SAP DNA binding domain, a central AAA+ domain, and a C-terminal RGG-type RNA-binding domain. SAF-A is important for restricting XIST RNA localization to the Xi chromosome territory during interphase, through a mechanism requiring the SAP and RGG domains. The presence of two spatially and functionally distinct nucleic acid binding domains suggested that SAF-A directly tethers XIST RNA to chromatin (Hasegawa et al., 2010). Subsequently, SAF-A has been implicated in a wide variety of nuclear RNA regulatory processes: forming interchromosomal connections through interactions with the FIRRE noncoding RNA (Hacisuleyman et al., 2014), mRNA splicing (Ye et al., 2015), and decompaction of euchromatic DNA through interaction with RNA (Nozawa et al., 2017). Notably, all described SAF-A functions occur in the interphase nucleus, and many of these processes are reversed during early mitosis. We therefore tested whether SAF-A is regulated during mitosis or if SAF-A-mediated chromosomal structures are remodeled during chromosome condensation.

In this work, we investigate the molecular mechanisms that regulate nuclear RNA localization during mitosis. We find that nuclear RNAs are removed from the surface of prophase chromosomes in an Aurora-B and SAF-A-dependent manner. We show that Aurora-B phosophorylates SAF-A at two sites in the SAP domain to release SAF-A:RNA complexes from chromatin during mitosis. Additionally, we find that nonphosphorylatable SAF-A phenocopies the genome-wide retention of RNA on mitotic chromosomes observed in Aurora-B inhibited cells, and leads to an increase in anaphase chromosome segregation defects. Our results show that removal of nuclear RNAs from chromatin is a key aspect of mitotic chromosomal restructuring and is essential for accurate chromosome segregation.

## Results

### SAF-A:RNA complexes undergo dynamic interactions with chromatin during the cell cycle

SAF-A is involved in several processes central to interphase nuclear function, including chromatin-bound RNA localization, interchromosomal interactions and DNA decondensation. Since these processes are all downregulated or reversed to allow for chromosome condensation and chromosome individualization during early mitosis, we hypothesized that SAF-A interactions with DNA or RNA were regulated in a cell cycle dependent manner.

To test for SAF-A:chromatin interactions, we monitored SAF-A localization across the cell cycle in human, diploid RPE-1 cells (Figure 1A). Interphase cells showed strong nuclear staining of SAF-A (Dreyfuss 1984; Hasegawa 2010), however SAF-A was dramatically cleared from chromatin in prophase, such that SAF-A staining of chromatin was absent by prometaphase. The exclusion of SAF-A from mitotic chromosomes was observed with two different antibodies to detect the native protein, in SAF-A-GFP transfected cells, and with mCherry knocked in to the endogenous SAF-A locus. To confirm that the exclusion of SAF-A from chromatin reflected a change in its physical association with chromatin during mitosis, we immunoprecipitated SAF-A in interphase and mitotic cell extracts and tested for the presence of the core histones in both control and α-SAF-A eluates (Figure 1B). Histone H3 was enriched in interphase α-SAF-A IPs relative to control IPs using IgG, but was absent in mitotic α-SAF-A IPs. Further, Coomassie-stained gel analysis of control and α-SAF-A IPs confirmed the presence of all four core histones in interphase but not mitotic α-SAF-A IPs. Together these data demonstrate SAF-A is removed from chromosomes in mitosis.

**Figure 1.**
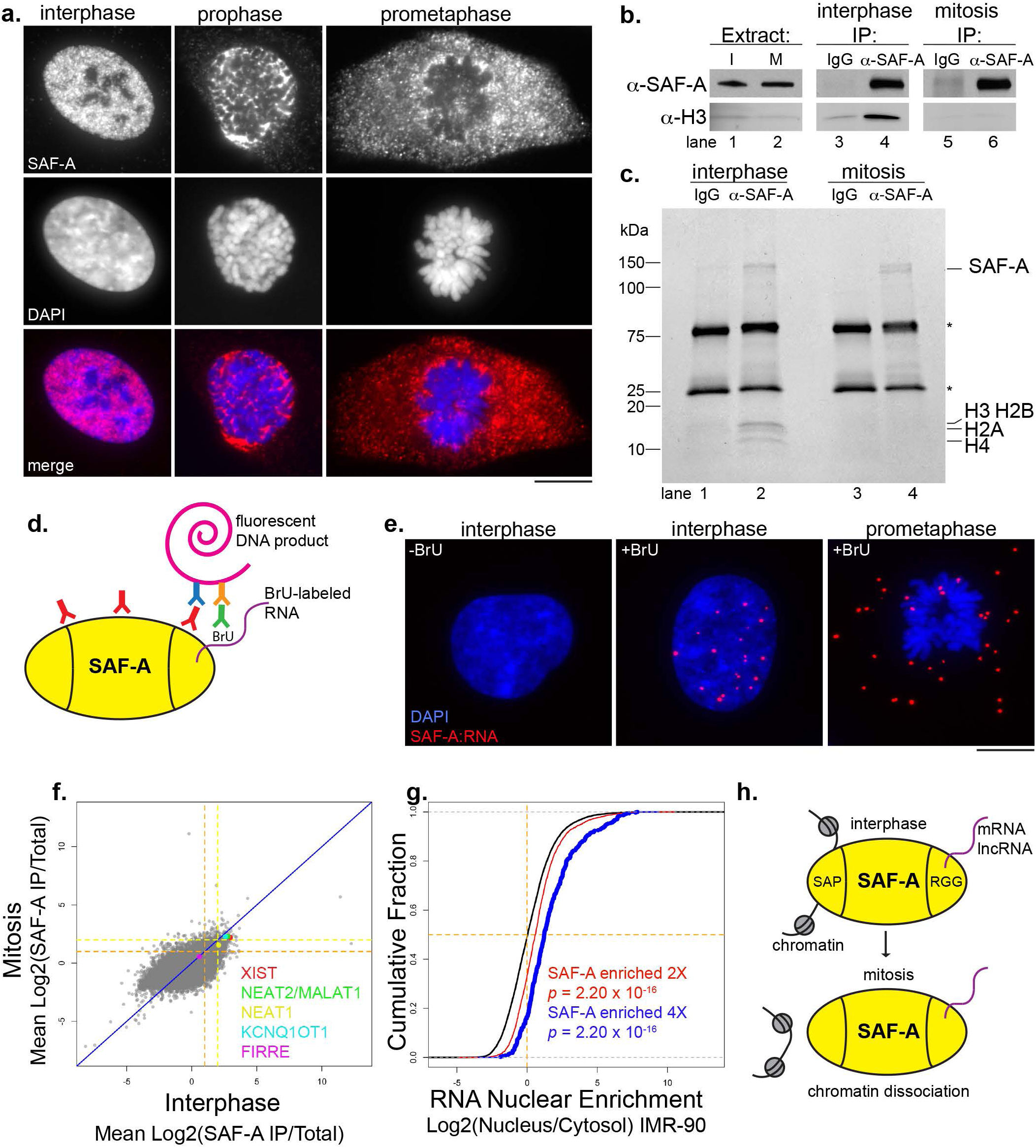
SAF-A:RNA complexes undergo dynamic interactions with chromatin during the cell cycle. **A.** Immunofluorescence of SAF-A in RPE-1 cells, relative to chromosomes (DAPI). Bar = 10 μm. **B.** Western blot analysis of SAF-A and histone H3 levels in cell extracts (lanes 1 and 2) and immunoprecipitations lanes 3-6) from cells arrested in interphase or mitosis. Mock IPs included mouse IgG (lanes 3 and 5); α-SAF-A IPs (lanes 4 and 6) were performed with a mouse monoclonal antibody specific for SAF-A. **C.** Coomassie-stained gel analysis of IgG IPs (lanes 1 and 3) and α-SAF-A IPs (lanes 2 and 4). IgG is denoted by the asterisks. **D.** The proximity ligation assay was modified to detect interactions between SAF-A and RNA. RPE-1 cells were labeled with BrU and incubated with primary antibodies to SAF-A and BrU. When the antigens are < 150 nm apart, secondary antibodies conjugated to oligonucleotides will generate a fluorescent DNA product. **E.** SAF-A:RNA PLA interactions detected in RPE-1 cells incubated with or without BrU to label RNA. **F.** RIP-seq was performed to identify SAF-A-interacting RNAs in interphase and mitotic RPE-1 cell extracts. To determine which RNAs associated with SAF-A in both interphase and mitosis, the average RNA enrichment in mitosis was plotted against RNA enrichment in interphase. Average enrichment values were determined from two independent biological replicates; all RNAs ≥ 2-fold enriched are gated by the dashed orange line, whereas all RNAs ≥ 4-fold enriched are gated by the dashed yellow line. **G.** Nuclear retention of SAF-A interacting RNAs. SAF-A interacting RNAs were compared to the nuclear/cytosolic distribution determined for each expressed RNA. SAF-A-interacting RNAs are shown in blue (4-fold enriched) and red (2-fold enriched); total RNA expression is shown in black. A rightward shift on the x-axis indicates a higher degree of nuclear retention for SAF-A-interacting RNAs relative to the total RNA population. **H.** Summary of SAF-A interactions with chromatin and RNA during interphase and mitosis.

Previous work has shown that SAF-A directly binds to hundreds of RNAs (Huelga et al., 2012; R. Xiao et al., 2012), but has not addressed whether SAF-A RNA binding is cell cycle-regulated. To determine whether differential chromatin association during the cell cycle influences SAF-A interactions with RNA, we modified the proximity ligation assay (PLA) to detect SAF-A:RNA complexes in asynchronous cell populations (Figure 1D). In this scheme, all RNA is labeled with BrdU prior to immunostaining with BrU and SAF-A primary antibodies. PLA detection of SAF-A:RNA interactions is then achieved through DNA polymerase amplification of oligonucleotide sequences conjugated to secondary antibodies. In interphase cells, SAF-A:RNA interactions were exclusively nuclear, whereas in mitotic cells, SAF-A:RNA interactions were dispersed throughout the entire cell (Figure 1E). Therefore, SAF-A maintains RNA interactions throughout the cell cycle, even after removal from chromosomes during mitosis, suggesting the two nucleic acid binding domains of SAF-A are regulated independently of each other.

To identify SAF-A-interacting RNAs across the cell cycle, we immunoaffinity purified SAF-A:RNA complexes from interphase and mitotic RPE-1 cell extracts and performed high-throughput sequencing (RNA-seq; Figure 1F). Sequencing reads from two independent biological replicates showed a high degree of correlation (Figure S1A-B), demonstrating the reproducibility of our sequencing data. Overall, we identified 1761 transcripts enriched ≥2-fold in SAF-A IPs from both interphase and mitotic cell extracts, representing 13% of all expressed RNAs. Quantitative RT-PCR confirmed the enrichment of specific transcripts in SAF-A IPs for all RNAs tested (Figure S1C). We noted several prominent lncRNAs such as XIST, NEAT2/MALAT1 and KCNQ1OT1 were enriched with SAF-A across the cell cycle, however, the majority of SAF-A-interacting transcripts were mature, fully spliced mRNAs (Figure S1C-D.

To determine if SAF-A-associated RNAs are preferentially retained in the nucleus, we compared our RNA-seq data with previous data examining relative nuclear enrichment of all RNAs in IMR-90 cells (Figures 1G and S1B; Djebali, 2012). Indeed, we found a highly statistically significant enrichment of nuclear-retained RNAs present in the SAF-A-interacting RNA population, and that a higher SAF-A RNA enrichment was correlated with a higher nucleus/cytosol localization ratio (Figure 1F, see 2-fold vs. 4-fold population). Taken together, our data show that SAF-A:RNA complexes are stable throughout the cell cycle, that SAF-A associates with hundreds of nuclear-retained mRNAs and lncRNAs during interphase, and that mitotic removal of SAF-A during prophase releases SAF-A:RNA from condensing chromosomes (Figure 1G).

### Aurora-B triggers relocalization of SAF-A:RNA complexes in early mitosis

Our data suggested there is a signal during prophase that triggers SAF-A removal from chromatin, prompting us to query whether inhibition of the Aurora-A, PLK1, or Aurora-B kinases that function during early mitosis would alter SAF-A localization in prometaphase. Whereas mitotic cells treated with DMSO showed the normal SAF-A chromatin exclusion pattern, treatment of cells with the Aurora-B selective inhibitor barasertib caused SAF-A enrichment on prometaphase chromosomes (Figure 2A, C). Similar results were observed using siRNAs to deplete Aurora-B (Figure 2B, C). In contrast, Plk1 or Aurora-A inhibition had no discernable effect on SAF-A localization (Figure 2C).

**Figure 2.**
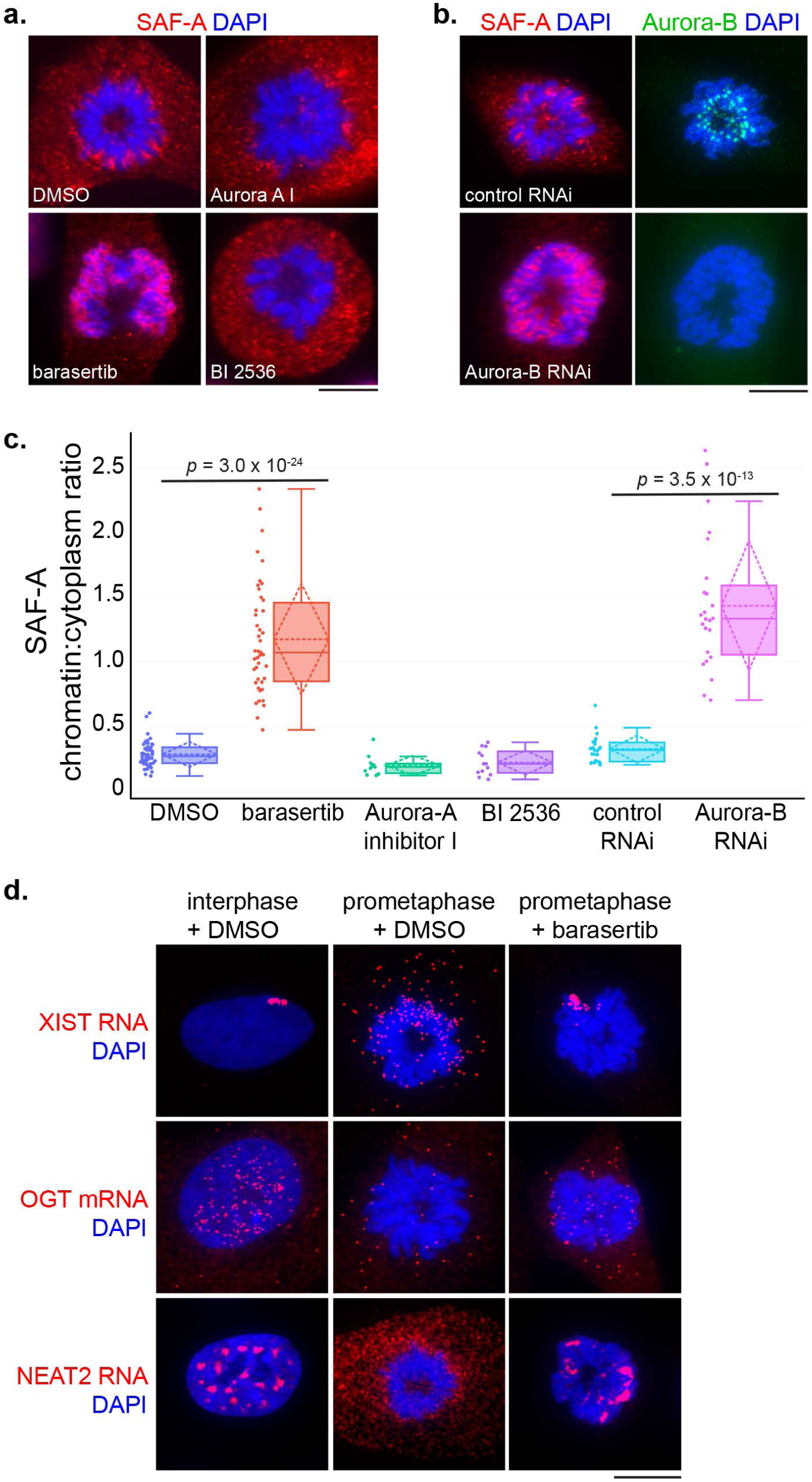
Aurora-B triggers relocalization of SAF-A:RNA complexes in early mitosis. **A.** SAF-A immunofluorescence in RPE-1 cells incubated with either DMSO or mitotic inhibitors to Aurora-B (barasertib), Aurora-A (Aurora-A inhibitor I), and Plk1 (BI 2536). Bar = 10 μm. **B.** RPE-1 cells were cultured in the presence of either control siRNAs or Aurora-B siRNAs. Cells were immunostained for SAF-A and Aurora-B. **C.** Image quantitation of mitotic SAF-A localization under the indicated treatments, expressed as a ratio of chromatin-localized SAF-A versus cytoplasmic SAF-A. For each boxplot in this study, the median is represented by the horizontal solid line within the shaded box; the mean and standard deviation are represented by the horizontal and diamond-shaped dashed line, respectively. Data points representing individual mitotic figures are rendered beside each boxplot in circles. DMSO, n = 49; barasertib, n = 43; Aurora-A inhibitor 1, n = 11; BI 2536, n = 13; control siRNA, n = 23; Aurora-B siRNA, n = 24. **D.** RNA FISH for the SAF-A-interacting XIST, OGT, and NEAT2 RNAs was performed in RPE-1 cells incubated with DMSO or barasertib.

To test whether Aurora-B inhibition caused SAF-A-interacting RNAs to be retained on chromatin, we treated cells with barasertib and performed FISH to detect three RNAs significantly enriched in SAF-A IPs: the XIST and NEAT2/MALAT1 lncRNAs and OGT mRNA (Figure 2D). In interphase, all three RNAs showed prominent nuclear retention, either as a chromosome-sized focus in the case of XIST (Brown et al., 1992), or multiple nuclear foci in the case of OGT and NEAT2. We note in contrast to the lncRNAs, OGT mRNA was also present as small granular cytoplasmic particles, consistent with it being a substrate for translation. In prometaphase cells treated with DMSO, XIST RNA was dispersed as small cytoplasmic foci; in contrast, prometaphase cells treated with barasertib showed an abnormal accumulation of XIST RNA on the mitotic Xi chromosome, consistent with previous observations (Clemson, McNeil, Willard, & Lawrence, 1996; Hall et al., 2009). A similar trend was observed for OGT and NEAT2 localization: whereas prometaphase cells showed a general trend of nuclear RNA disaggregation and cytoplasmic dispersal, Aurora-B inhibition resulted in markedly increased overlap of OGT and NEAT2 RNAs with mitotic chromosomes. Together, these data suggest the Aurora-B kinase is responsible for the removal of SAF-A:RNA complexes from prophase chromatin.

To determine whether Aurora-B has a global role in regulating RNA localization, we labeled cells with a 3 hour pulse of 5-ethynyl uridine (EU) (Jao & Salic, 2008) and monitored total RNA localization in prometaphase cells with or without Aurora-B inhibition (Figure 3). In control cell populations, we observed that EU-labeled RNA was substantially enriched in the nucleus relative to the cytoplasm, as previously observed (Figure 3A, C)(Jao & Salic, 2008; Johnson et al., 2017; Palozola et al., 2017). However, RNA was excluded from condensed prophase and prometaphase chromosomes (Figure 3B). Strikingly, barasertib-treated cells showed RNA retention on the chromosome surface, reminiscent of the SAF-A localization pattern in Aurora-B-inhibited cells (Figure 3B, D). Therefore, Aurora-B releases a large population of nuclear/chromosomal RNAs from chromosomes in prometaphase.

**Figure 3.**
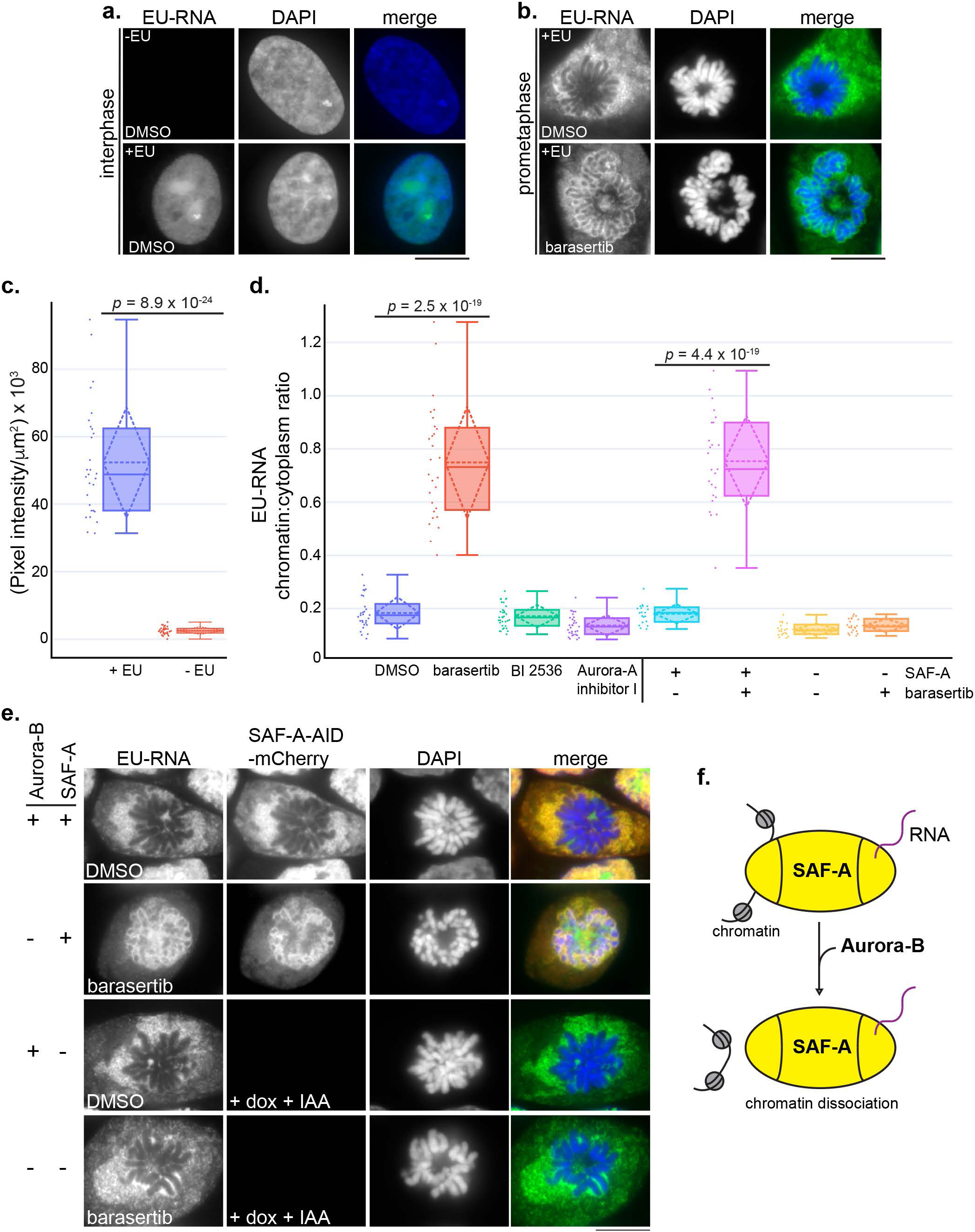
Aurora-B and SAF-A together define a pathway for removing RNP complexes from mitotic chromosomes. **A-B.** EU-labeled RNA was detected in interphase and prometaphase DLD-1 cells treated with DMSO or barasertib. Chromosomes were stained with DAPI. Bar = 10 μm. **C.** Quantitative image analysis demonstrated a 21-fold increase in fluorescence intensity measured in + EU-labeled cell populations (n = 29) relative to control cells (n = 31). **D.** Quantitation of mitotic EU-RNA localization, expressed as a ratio of chromatin-localized RNA versus cytoplasmic RNA. Cells were treated with DMSO or mitotic inhibitors as shown. DMSO, n = 29; barasertib, n = 28; BI 2536, n = 34; Aurora-A inhibitor 1, n = 31. **E.** EU-RNA was detected in DLD-1 cells homozygous for a SAF-A-AID-mCherry degron allele. Cells were treated with either DMSO or barasertib, with or without SAF-A depletion (+dox +IAA), to determine the epistasis relationship of Aurora-B and SAF-A. Quantitation of mitotic EU-RNA localization in SAF-A degron cells under all conditions is shown in **C.** SAF-A undepleted + DMSO n = 20; SAF-A undepleted + barasertib n = 25; SAF-A depleted + DMSO n = 20; SAF-A depleted + barasertib n = 22. **F.** Model for Aurora-B dependent regulation of SAF-A:RNA chromatin association.

If Aurora-B primarily targets SAF-A to remove RNAs from chromatin in mitosis, we hypothesized that SAF-A ablation should result in RNA release from chromatin in Aurora-B-inhibited cells. To test this, we constructed a human diploid DLD-1 cell line with both copies of endogenous SAF-A fused to an auxin-inducible degron sequence and mCherry (SAF-A-AID-mCherry; Figure S2A-B) (Holland, Fachinetti, Han, & Cleveland, 2012; Natsume, Kiyomitsu, Saga, & Kanemaki, 2016); a doxycycline-inducible TIR1 E3 ligase was also integrated to enable auxin-mediated degradation of SAF-A-AID-mCherry. Control experiments demonstrated that SAF-A-AID-mCherry was homozygous, expressed at normal levels, showed a localization pattern identical to the wild-type protein, and could be depleted within 24 hours of treatment with doxycycline and auxin (Figure S2C-D). We then tested for EU-RNA localization under all combinations of SAF-A-AID-mCherry depletion and Aurora-B inhibition. In cells expressing SAF-A-AID-mCherry, EU-RNA was retained on chromatin in the presence of barasertib, as was observed in wild-type cells (Figure 3E). However, in SAF-A-depleted cells, Aurora-B inhibition no longer caused EU-RNA retention on chromosomes, demonstrating that Aurora-B is epistatic to SAF-A (Figure 3D-E). Collectively, we conclude Aurora-B and SAF-A together define a pathway that releases RNA from chromosomes *en masse* during early mitosis (Figure 3F).

### Phosphorylation of SAF-A by Aurora-B controls chromatin association

To test whether Aurora-B promotes chromosomal removal of SAF-A through direct phosphorylation, we performed a kinase assay using recombinant Aurora B:Incenp complex (Bolton et al., 2002; Rosasco-Nitcher, Lan, Khorasanizadeh, & Stukenberg, 2008) and immunoaffinity-captured SAF-A (Figure 4A). We observed dose-dependent phosphorylation of SAF-A in the presence of the active Aurora B-Incenp complex. A low level of SAF-A phosphorylation was present in reactions lacking Aurora B-Incenp, possibly due to a trace amount of kinase activity copurifying with SAF-A (Figure 3A, lane 5). In contrast, there were no specific phosphorylation events observed in IgG control immunoprecipitations. Therefore, the Aurora B-Incenp complex can directly phosphorylate SAF-A.

**Figure 4.**
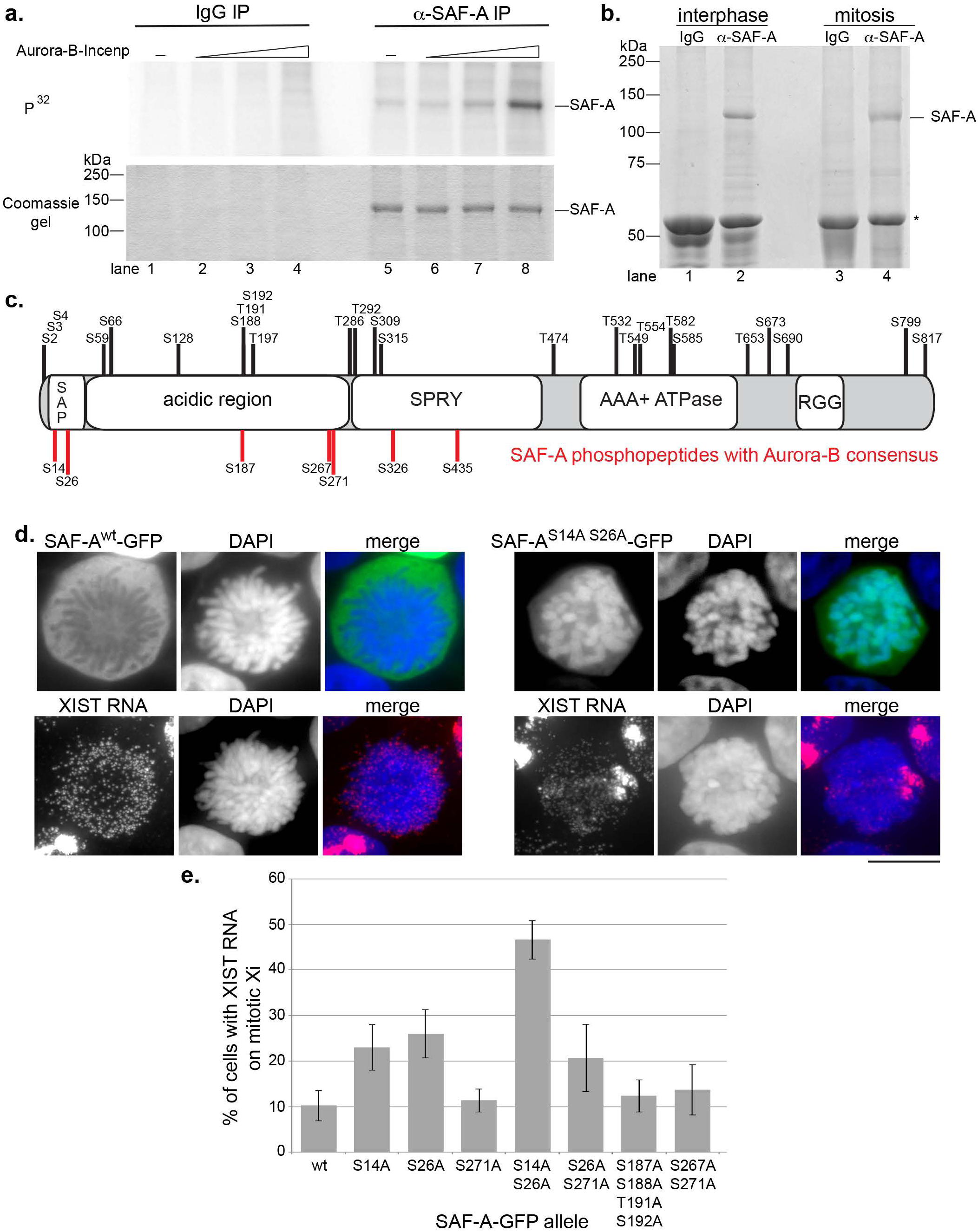
Aurora-B phosphorylates SAF-A on residues S14 and S26 of the SAP domain. **A.** IP kinase assay. Control IgG IPs (lanes 1-4) and α-SAF-A IPs (lanes 5-8) were incubated with purified Aurora-B:Incenp complex. Phosphorylation of SAF-A was detected using ^32^P-labeled ATP (upper panel), and SAF-A was detected using coomassie blue (lower panel). Aurora-B-Incenp was present at 0 nM (lanes 1 and 5), 1 nM (lanes 2 and 6), 10 nM (lanes 3 and 7), or 100 nM (lanes 4 and 8). **B.** SAF-A was immunopurified from interphase and mitotic cell extracts for mass spectrometry identification of phosphopeptides. IgG heavy chain is denoted by the asterisk. **C.** Phosphorylated serines and threonines were mapped relative to the domain structure of SAF-A as described in the text. Serines depicted in red fit the Aurora-B consensus sequence (R/K-X-S/T, or R/K-X-S/T); serines and threonines depicted in black do not have the Aurora-B consensus motif. **D.** GFP immunofluorescence for wild-type and phosphomutant versions of SAF-A -GFP expressed in HEK293T cells, combined with XIST RNA FISH. Bar = 10 μm. **E.** Quantitation of the percent of cells expressing either wild-type or phosphomutant SAF-A-GFP showing retention of XIST RNA on the mitotic Xi. 100 cells were scored for each transfection; the average and standard deviation of multiple independent transfections is shown. For SAF-A^wt^-GFP and SAF-A^S14A S26A^-GFP n = 5; n = 3 for all other alleles tested.

To identify phosphorylation sites on SAF-A *in vivo*, we isolated SAF-A from interphase and mitotic cell extracts and used mass spectrometry to identify modified peptides (Figure 4B; Supplemental Table 1). We pooled our phosphopeptide data with that from several phosphoproteomic studies (www.phosphosite.org) and mapped serine and threonine phosphorylation sites relative to SAF-A domain structure (Figure 4C). Potential Aurora-B sites were then identified based on whether the proximal amino acids fit the Aurora-B consensus sequence (K/R-S/T or K/R-X-S/T(Alexander et al., 2011; Cheeseman et al., 2002; Hengeveld et al., 2012; Kettenbach et al., 2011)). All seven of the putative Aurora-B sites were located in the N-terminal half of SAF-A, with two positioned in the SAP domain required for DNA binding (S14, S26), three positioned in a low complexity acidic domain (S187, S267, S271), and two located within the SPRY domain (S326, S435). Phosphorylated SAF-A S14, S26, S59, S187, and S271 were overrepresented in mitosis (Supplemental Table 1;(Douglas et al., 2015; Kettenbach et al., 2011; Olsen et al., 2010)).

To determine if the predicted Aurora-B phosphorylation sites regulate SAF-A:RNA localization during mitosis, we constructed a series of phosphomutants with single or multiple alanine substitutions in the SAP domain and acidic domain, and analyzed mitotic 293T cells expressing wild-type or phosphomutant SAF-A-GFP for XIST RNA localization (Figure 4D-E). Cells expressing wild-type SAF-A-GFP showed normal localization of the tagged protein and XIST RNA: both components showed nuclear localization during interphase and exclusion from mitotic chromosomes. In contrast, we found that mutation of S14A and S26A, either alone or in combination, significantly increased the number of cells showing ectopic retention of SAF-A on mitotic chromosomes and focal XIST RNA staining on the Xi during prometaphase (Figure 4D). In particular, prometaphase cells expressing SAF-A^S14A S26A^-GFP had a 50% frequency of cells with focal, chromatin-bound XIST RNA staining, nearly a 5-fold increase relative to the SAF-A^wt^-GFP control (Figure 4E). Expression of either SAF-A^S14A^-GFP or SAF-A^S26A^-GFP caused a less dramatic yet significant effect, with both causing a ~2-fold increase in prometaphase cells with focal XIST localization. Further, cells expressing SAF-A^S26A S271A^-GFP had an XIST RNA localization pattern similar to SAF-A^S26A^-GFP, demonstrating that the cumulative effects of mutating two phosphorylatable serines was specific to the SAF-A SAP domain residues S14 and S26. The other SAF-A phosphomutants analyzed had no effect on XIST RNA localization distinguishable from SAF-A^wt^-GFP. Further, an alignment of SAF-A from multiple species showed conservation of S14 and S26 residues throughout the vertebrate lineage (Figure S3). Based on these data we conclude that S14 and S26 regulate mitotic SAF-A localization.

### Phosphorylation of the SAP domain reduces SAF-A binding to DNA

Previous work has demonstrated that the SAF-A can bind directly to AT-rich DNA *in vitro*, and that the N-terminal SAP domain is a DNA-binding domain (Fackelmayer, Dahm, Renz, Ramsperger, & Richter, 1994; Gohring, Schwab, Nicotera, Leist, & Fackelmayer, 1997; Kipp et al., 2000; Romig, Fackelmayer, Renz, Ramsperger, & Richter, 1992). To gain molecular insight as to how phosphorylation of residues S14 and S26 on SAF-A could block DNA binding, we performed molecular modeling of the SAF-A SAP domain using I-TASSER software, which uses a combination of *ab initio* modeling and refinement based on known structures in the protein data bank (Figure 5A; (Yang et al., 2015). I-TASSER identified several close sequence and structural homologs of SAF-A, all of which featured a SAP domain with a helix-turn-helix structural motif. Using this model we found that S14 is positioned within the first helix, whereas S26 is positioned in a loop between the first and second helix (depicted in yellow, Figure 5A).

**Figure 5.**
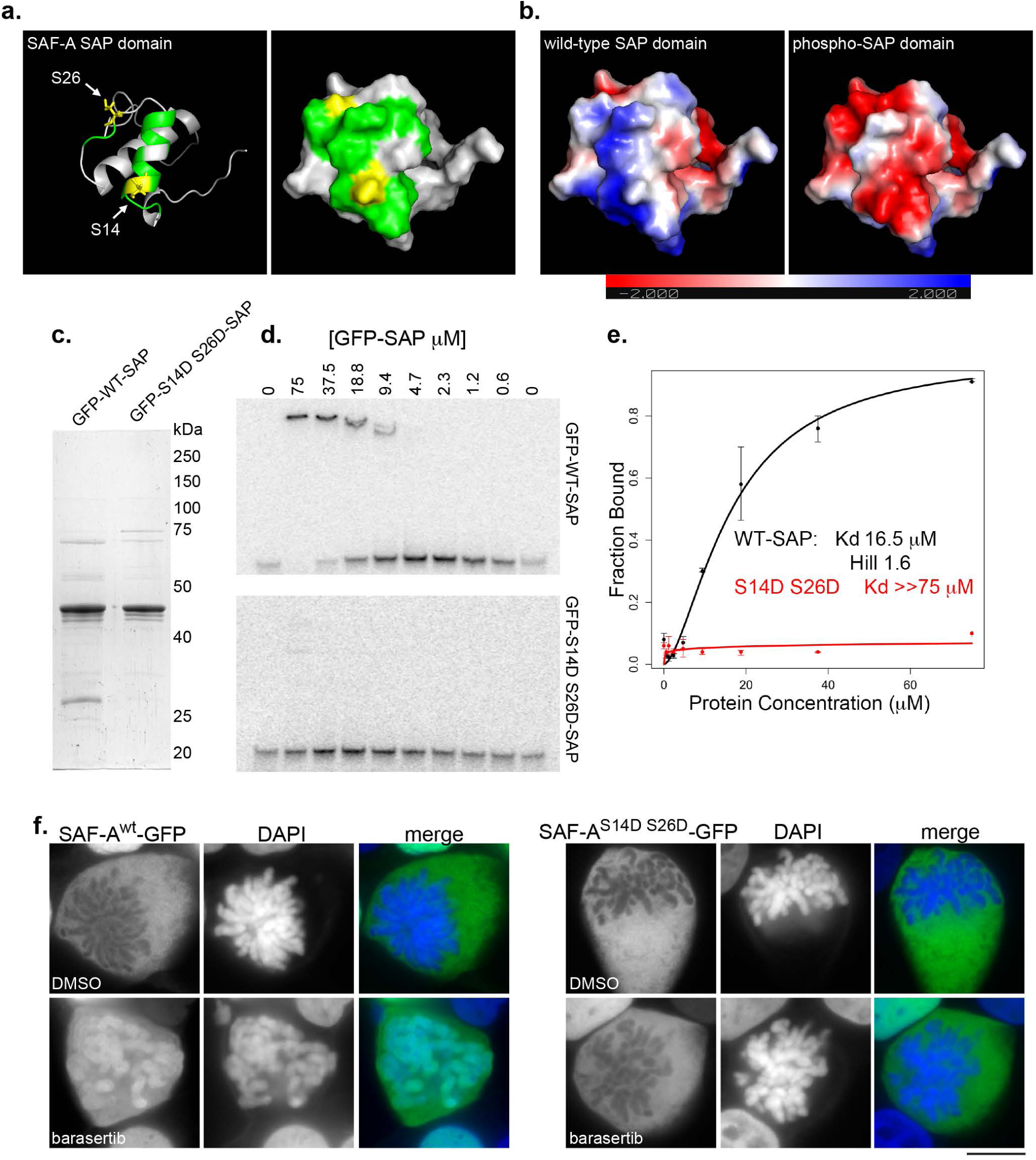
Aurora-B phosphorylation of the SAP domain reduces SAF-A binding to DNA. **A.** Molecular modeling of the SAP domain of SAF-A, rendered as a ribbon model (left), and as a space-filling model (right). Residues S14 and S26 are highlighted in in yellow; residues predicted to define the DNA-binding interface are colored in green. **B.** Electrostatic surface charge potential was determined for the unphosphorylated SAP domain (left) or the SAP domain phosphorylated on residues S14 and S26 (right). The color gradient inset shows the charge distribution as a continuum between red (more negative) and blue (more positive). **C.** Purification of recombinant wild-type (GFP-WT-SAP) and phosphomimetic (GFP-S14D S26D-SAP) SAP domains of SAF-A from *E. coli*. **D.** DNA electrophoretic mobility shift assays (EMSA) for GFP-WT-SAP and GFP-S14D S26D-SAP with an AT-rich DNA template. **E.** The fraction of bound DNA template versus the protein concentration of GFP-WT-SAP or GFP-S14D S26D-SAP was plotted to determine the dissociation constant, K_d_, and the Hill coefficient. Error bars are standard deviation of two replicates. **F.** Wild-type and phosphomimetic alleles of SAF-A-GFP were expressed in HEK293T cells in the presence of either DMSO or barasertib. SAF-A^wt^-GFP and SAF-A^S14D S26D^-GFP were visualized by GFP immunostaining. Bar = 10 μm.

We used the NMR structure of the SAP domain-containing protein PIAS1 to map the residues contacting DNA onto the equivalent positions in the SAF-A model (depicted in green; Figure 5A)(Okubo et al., 2004), and observed both S14 and S26 were positioned on the same molecular surface of the SAP domain as the residues predicted to contact DNA. To test whether phosphorylation of residues S14 and S26 altered the electrostatic surface potential of the SAP domain, we calculated the surface charge for both the unphosphorylated and phospho-S14 S26 models of the SAF-A SAP domain. We found that the unphosphorylated SAF-A SAP domain has an overall net positive charge (Figure 5B, blue), and that this charge is dramatically reversed by phosphorylation of S14 and S26 (Figure 5B, red). These data suggest that charge repulsion between the phospho-SAP domain and the DNA phosphate backbone is a likely mechanism for the release of SAF-A from chromatin after phosphorylation by Aurora-B.

To directly test if altered surface charge potential of the SAF-A SAP domain affects DNA binding, we reconstituted DNA binding by the SAF-A SAP domain *in vitro* using GFP-SAP^WT^ and GFP-SAP^S14D S26D^ expressed and purified from *E. coli* (Figure 5C). We then employed a DNA EMSA using a 50 bp AT-rich SAR sequence (Okubo et al., 2004) to measure the DNA binding affinity of each protein (Figure 5D-E). SAP^WT^ domain readily bound the SAR DNA with an apparent K_d_ of 16.5 μM. In contrast, the SAP^S14D S16D^ domain displayed a dramatically reduced DNA binding, such that we were only able to detect minimal DNA binding activity at the highest protein concentration tested (K_d_>>75 μM). We conclude that phosphorylation of the SAP domain of SAF-A changes the surface charge of the DNA binding domain and causes dramatically reduced DNA binding.

The reduced DNA binding affinity of the phosphomimetic SAP domain *in vitro* suggested that a phosphomimetic SAF-A allele would show reduced chromatin affinity *in vivo*. We therefore compared the localization of SAF-A^wt^-GFP to SAF-A^S14D S26D^-GFP, with and without barasertib treatment (Figure 5F). In barasertib-treated cell populations, 95% of cells showed SAF-A-wt-GFP retained on prometaphase chromosomes compared to only 3% of cells treated with DMSO. In contrast, the localization of SAF-A^S14D S26D^-GFP was completely excluded from mitotic chromosomes with or without barasertib treatment, thus demonstrating the Aurora-B inhibitor acts through SAF-A S14 and S26. Taken together, our mutant analysis data indicate that Aurora-B phosphorylates SAF-A on residues S14 and S26 in the DNA binding-SAP domain *in vivo*, and that phosphorylation of these two residues acts additively to dissociate SAF-A:RNA complexes from chromatin during early mitosis.

### Evidence that monomeric SAF-A mediates RNA:DNA tethering

The domain structure of SAF-A includes a central AAA^+^-type ATPase domain that controls protein oligomerization through ATP binding and hydrolysis (Nozawa et al., 2017). Consequently, mutations in specific Walker box residues can be used to manipulate the SAF-A oligomeric state (Figure 6A). SAF-A exists as a monomer in its most abundant cellular form, yet it has not been determined whether the monomeric or oligomeric form of SAF-A contributes to RNA localization. XIST RNA FISH in mitotic cells expressing the SAF-A phosphomutant allowed direct visualization of SAF-A:RNA:chromatin interactions *in vivo*. To determine whether SAF-A mediates XIST RNA localization as a monomer or SAF-A oligomer we generated SAF-A^K510A^-GFP (monomeric) and SAF-A^D580A^-GFP (oligomeric) alleles, either alone or in combination with the SAF-A^S14A S26A^-GFP phosphomutant, and monitored mitotic XIST RNA localization in transfected cell populations (Figure 6B). Interestingly, we found that the monomeric SAF-A phosphomutant (SAF-A^S14 S26 K510A^-GFP) anchored XIST RNA to mitotic chromatin nearly as efficiently as the SAF-A^S14 S26^-GFP allele. In contrast, the oligomeric SAF-A phosphomutant (SAF-A^S14 S26 D580A^) was indistinguishable from the wild-type allele. These data suggested that the monomeric form of SAF-A is sufficient to tether RNA to chromosomes.

**Figure 6.**
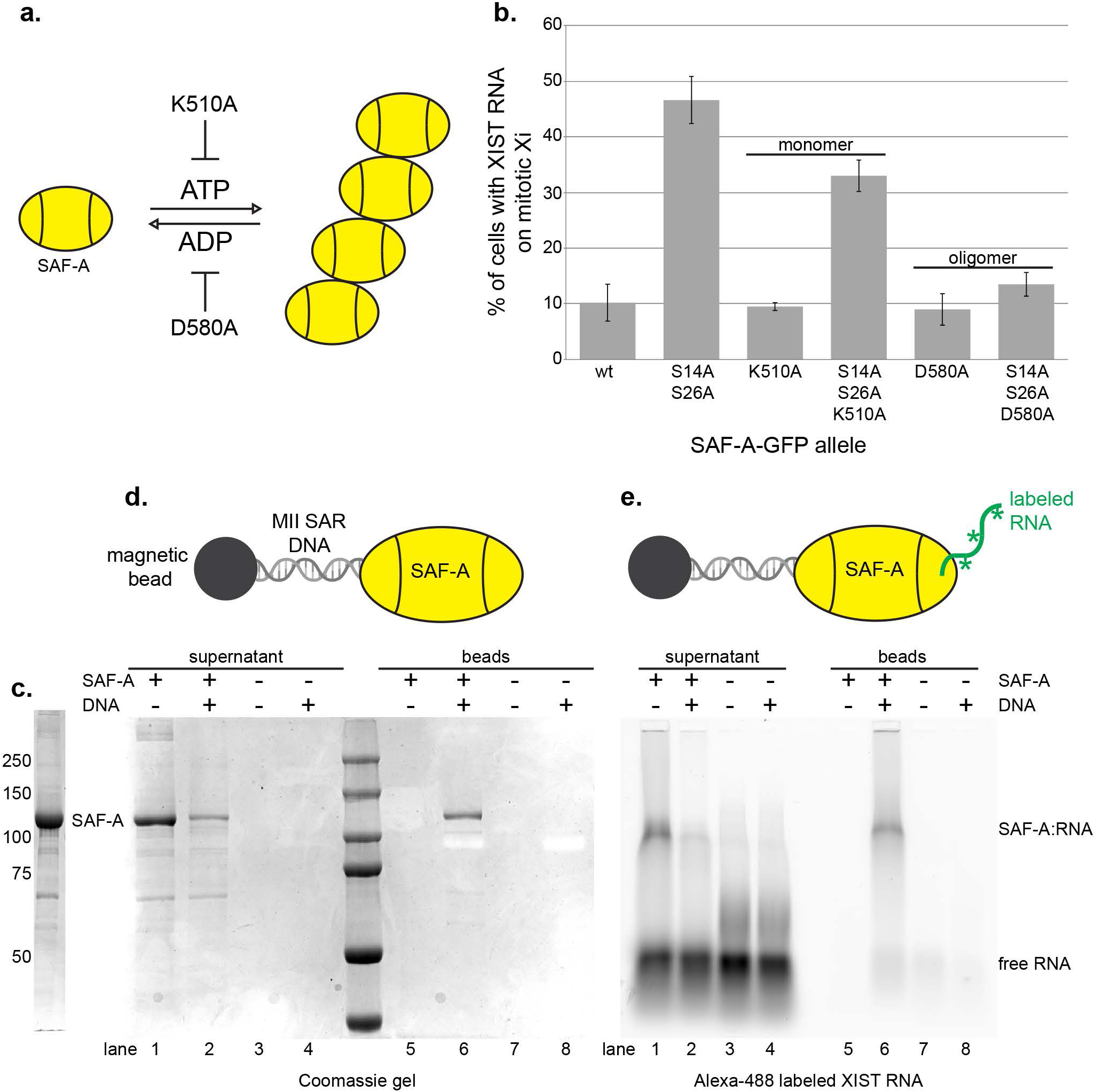
Monomeric SAF-A is sufficient for RNA tethering. **A.** Model of ATP-dependent oligomerization cycle of AAA^+^-type ATPases. The K510A mutation in the SAF-A Walker A box inhibits ATP binding and protein oligomerization, while the D580A mutation in the Walker B box inhibits ATP hydrolysis, trapping SAF-A in the oligomeric state. **B.** Quantitation of percent of cells showing XIST RNA retention on the mitotic Xi in HEK293T cells transfected with SAF-A -GFP alleles. 100 cells were scored for each transfection; the average and standard deviation of multiple independent transfections is shown. For SAF-A^wt^-GFP and SAF-A^S14A S26A^-GFP n = 5; n = 2 for all other alleles tested. **C.** Purification of full-length, monomeric SAF-A from Sf9 cells. **D.** Capture of SAF-A on DNA coated beads. The SAF-A-interacting DNA sequence from the MII SAR element was conjugated to magnetic beads, and used to capture SAF-A. Coomassie gel staining was used to examine the presence of unbound SAF-A in reaction supernatants (lanes 1-4), and SAF-A binding to DNA coated beads (lanes 5-8). **E.** RNA tethering assay. SAF-A complexed with an FITC labeled XIST RNA fragment (lanes 5-8) was incubated with DNA beads. FITC fluorescence was used to image SAF-A:RNA complexes in reaction supernatants (lanes 1-4), and retention of RNA with SAF-A bound to DNA beads (lanes 5-8).

To test this idea, we devised an *in vitro* RNA:DNA tethering assay and queried whether monomeric SAF-A could bridge interactions between DNA-conjugated beads and RNA. We expressed and purified full-length monomeric SAF-A protein (Figure 6C), and used a nuclear scaffold DNA sequence coupled to magnetic beads to capture SAF-A in the absence of ATP to block polymerization (MII SAR DNA; (Fackelmayer et al., 1994)). SAF-A bound efficiently to DNA-conjugated beads, without detectable binding to control beads (Figure 6D). To assay whether SAF-A bound to DNA could simultaneously bind RNA, we incubated SAF-A complexed with a fragment of the XIST RNA with DNA beads (Huelga et al., 2012), and found that XIST RNA was retained specifically on SAF-A:DNA beads (Figure 6E). These data support the idea that SAF-A has intrinsic RNA:DNA tethering activity in the absence of protein oligomerization or accessory proteins. Given the abundance of SAF-A is estimated to be 2 × 10^6^ molecules per cell (Fackelmayer & Richter, 1994), our data suggest numerous DNA:SAF-A:RNA bridges are formed in the interphase nucleus.

### Dynamic localization of SAF-A promotes normal chromosome segregation in mitosis

The data suggested that removal of SAF-A:RNA complexes from chromatin is a normal feature of prometaphase, prompting us to test whether the role of Aurora-B in determining mitotic localization of SAF-A is important for normal chromosome transmission. For this purpose, we used lentiviruses to integrate tet-inducible versions of SAF-A-GFP or SAF-A-^S14A S26A^-GFP into the human cell line engineered with the tagged, degradable SAF-A-AID-mCherry (Figure 7A). To simultaneously deplete SAF-A-AID-mCherry and reconstitute SAF-A function with the GFP-tagged alleles, cells were treated with doxycycline for 24 hours to induce SAF-A-GFP and TIR1, and then subsequently treated with both doxycycline and auxin for another 24 hours to induce degradation of SAF-A-AID-mCherry prior to analysis (Figure 7B).

**Figure 7.**
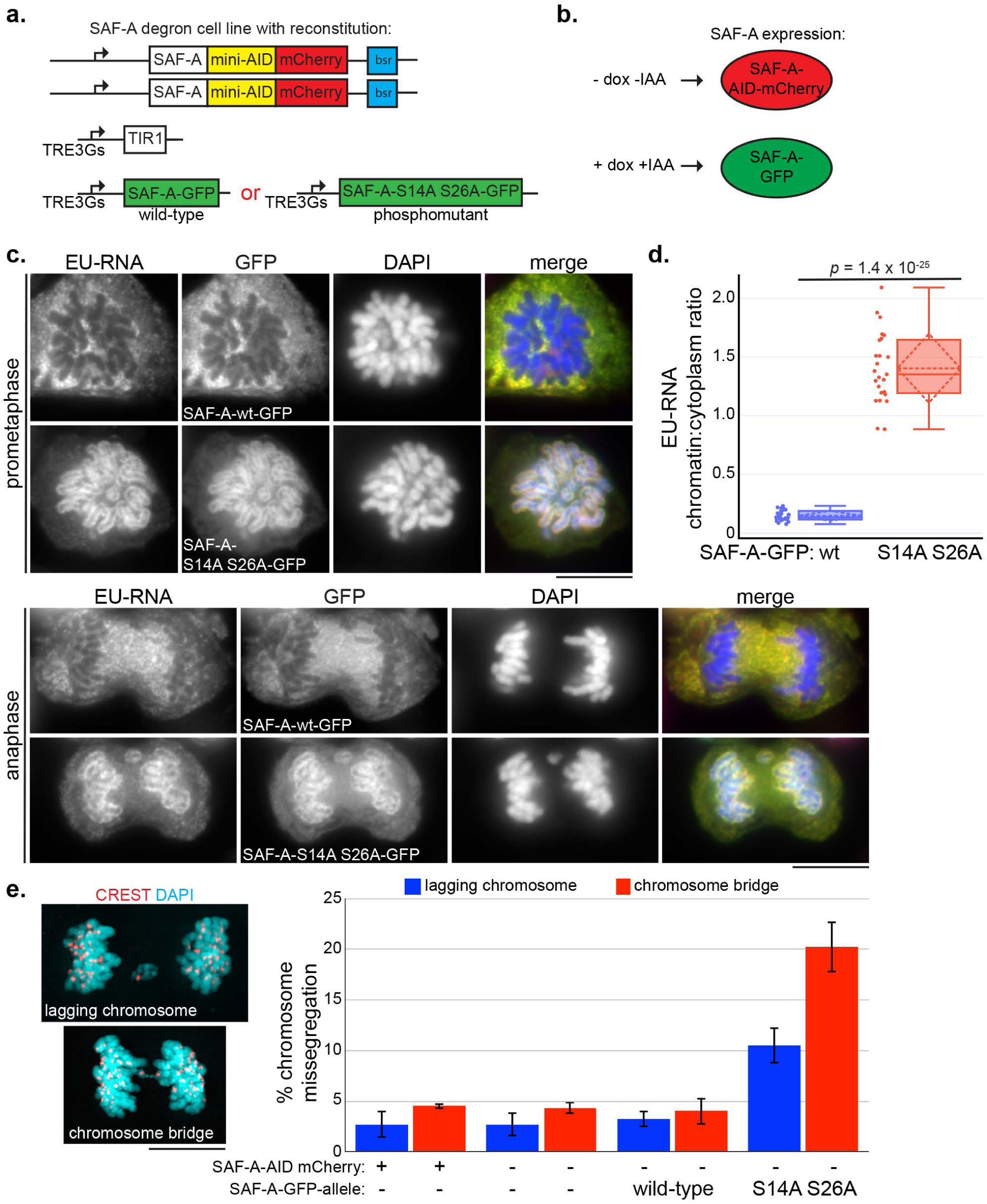
Dynamic localization of SAF-A promotes normal chromosome segregation in anaphase. **A.** SAF-A-AID-mCherry degron cells were transduced with lentivirus encoding either SAF-A-wt-GFP or the phosphomutant SAF-A^S14A-S26A^-GFP. **B.** In this scheme, untreated cells express only SAF-A-AID-mCherry. Addition of doxycycline and auxin (IAA) induces SAF-A-GFP alleles while also causing degradation of SAF-A-AID-mCherry protein through induction of TIR1. **C.** The SAF-A degron cell line was reconstituted with either SAF-A^wt^-GFP or SAF-A^S14 S26^-GFP. Cells were labeled with EU to detect RNA localization and immunostained for GFP to detect SAF-A-GFP alleles. Examples of cells in prometaphase and late anaphase are shown. **D.** Quantitation of EU-RNA localization in SAF-A-GFP-reconstituted cell lines, expressed as a ratio of chromatin-associated RNA versus cytoplasmic RNA. SAF-A^wt^-GFP n = 25; SAF-A^S14A S26A^ n = 25. **E.** Chromosome missegregation during anaphase was measured in cells with and without SAF-A-AID-mCherry depletion, and in cells reconstituted with SAF-A^wt^-GFP or SAF-A^S14 S26^-GFP. Three biological replicates were performed, with 200 anaphases scored in each experiment for the incidence of lagging chromosomes and chromosome bridge formation. Error bars indicate standard deviation. An example of each type of aberrant anaphase is depicted beside the graph with CREST immunostaining to mark kinetochores. Bar = 10 μm.

To determine whether the GFP-tagged SAF-A alleles showed localization properties predicted by our data, we visualized GFP immunofluorescence together with RNA localization (Figure 7C). Inspection of individual mitotic figures revealed that the SAF-A-GFP cell line showed exclusion of the protein and EU-RNA from chromosomes, comparable to endogenous RNA and SAF-A localization in untreated wild-type cells. Conversely, cells expressing the unphosphorylatable mutant SAF-A-^S14A S26A^-GFP showed retention of SAF-A and RNA on chromosomes, in a pattern that persisted throughout mitosis. Quantitation of EU-RNA confirmed these observations, as cells expressing the SAF-A phosphomutant had 9.6-fold more chromosome-associated RNA than SAF-A-wt-GFP cells (Figure 7D). These data confirm that the SAF-A allele reconstitution assay recapitulates the phenotypes predicted by chemical inhibition data (Figures 2–3) and transient transfection (Figure 4). Further, the data strongly argue that Aurora-B causes relocalization of nuclear and chromosomal RNAs during mitosis through targeting S14 and S26 in the SAF-A SAP domain.

To test whether mitotic RNA localization impacts chromosome inheritance in anaphase, we stained SAF-A-GFP and SAF-A-^S14A S26A^-GFP cells with a CREST antibody that labels kinetochores, and scored anaphases in asynchronous cell populations (Figure 7E). We observed that wild-type DLD-1, undepleted SAF-A-AID-mCherry cells, depleted SAF-A-AID-mCherry cells, and cells expressing SAF-A-GFP all displayed a comparable low frequency of aberrant anaphases. In contrast, cells expressing SAF-A-^S14A S26A^-GFP showed a significant increase in the rate of lagging chromosomes present at the midzone (3.3-fold increase; *p* = 2.9 × 10^−3^) and the rate of anaphase chromosome bridge formation (5-fold increase; *p* = 6.0 × 10^−4^). We conclude that the proper mitotic localization of SAF-A:RNA complexes is important for normal chromosome segregation in anaphase.

## Discussion

In this work, we show that Aurora-B phosphorylates the SAF-A SAP domain to remove nuclear RNAs from the surface of all chromosomes during prophase. Further, we demonstrate that removal of mitotic SAF-A:RNP complexes from the chromosome surface is important for accurate chromosome segregation during anaphase. Our work identifies a previously unrecognized chromosomal remodeling process that is temporally correlated with other genome restructuring events during early mitosis.

During prophase, chromosome structure is dramatically remodeled through the coordinated action of several pathways that impact chromosome condensation, sister chromatid resolution, and transcriptional silencing. Recent work using Hi-C methods demonstrated that interphase chromosomal structures such Topologically Associated Domains (TADs) and A/B compartments are removed from chromosomes during the first 15 minutes of mitosis, and are replaced by a structure of nested loop domains orchestrated by the condensin I and II complexes (Gibcus et al., 2018; Naumova et al., 2013; Walther et al., 2018). During interphase, TADs are maintained by the combined action of the cohesin complex and CTCF, which together facilitate interactions between enhancer and promoter regions (Kagey et al., 2010; Nora et al., 2017; Rao et al., 2017). Quantitative proteomics of chromosomal proteins during mitosis showed that the cohesin complex is removed from chromosome arms at the same time as condensin complexes begin to associate with chromosomes (Gibcus et al., 2018; Ohta et al., 2010). Mitotic transcriptional silencing is synchronous with mitotic chromosomal remodeling, and occurs through the phosphorylation of transcription initiation factors and run-off of elongating RNAPII (Akoulitchev & Reinberg, 1998; Liang et al., 2015; Segil et al., 1996). Our work shows that in addition to these chromosome remodeling events, chromosome-associated RNP complexes are removed during chromosome condensation. Thus, during prophase multiple different pathways function in parallel to erase interphase genome structure and condense chromosomes in preparation for chromosome segregation.

In early mitosis, Aurora-B regulates chromosome structure through cohesin removal, condensin I loading, and heterochromatin dissociation (Fischle et al., 2005; Giet & Glover, 2001; Hirota et al., 2005; Lipp, Hirota, Poser, & Peters, 2007; Losada, Hirano, & Hirano, 2002). We now show that phosphorylation of SAF-A by Aurora-B also contributes to chromosomal remodeling by releasing chromatin-bound RNAs. Interestingly, Aurora-B phosphorylates the pluripotency transcription factor Oct4 in ES cells to promote chromatin release during mitosis (Shin et al., 2016). Collectively, these observations demonstrate that Aurora-B controls many different pathways important for restructuring interphase chromatin during early mitosis. The fact that Aurora-B triggers the release of chromatin-bound RNAs and core transcription factors suggests that Aurora-B may be a key factor responsible for resetting the transcriptional program as cells pass through mitosis. Passage through mitosis is a key step mediating transcriptional reprogramming and cell fate transitions (Egli, Birkhoff, & Eggan, 2008; Soufi & Dalton, 2016), and our work suggests that phosphorylation of SAF-A may be a key component of this process.

SAF-A has been implicated in several processes that control interphase genome structure, all of which are mediated by SAF-A interactions with RNA. For example, SAF-A impacts inactive X chromosome structure through the control of XIST RNA:Xi localization (Hasegawa et al., 2010; Helbig & Fackelmayer, 2003; Pullirsch et al., 2010) SAF-A is also required for nuclear localization of the FIRRE RNA, which promotes interchromosomal interactions between a subset of autosomes (Hacisuleyman et al., 2014). In this study we identify approximately 1800 additional RNAs that associate with SAF-A throughout the cell cycle, therefore significantly expanding the known repertoire of known SAF-A:RNA interactions. Further, because we could visualize SAF-A:RNA complexes bound to chromosomes in total RNA labeling experiments, we hypothesize that many of the SAF-A:RNA complexes we have identified are anchored to chromatin in a similar fashion as the XIST and FIRRE RNAs. Indeed, based on cytological evidence (Figure 3) it appears that most or all of Aurora-B regulated chromatin-associated RNA localization depends on SAF-A function.

In addition to its role in RNA localization, SAF-A promotes decondensation of expressed regions of the interphase genome by interacting with nuclear RNAs (Fan et al., 2018; Nozawa et al., 2017). Interestingly, the decondensation function of SAF-A is linked to SAF-A oligomerization through the AAA+ domain and RNA-binding activity, but is independent of the DNA binding domain. Since this study did not explicitly address RNA localization, it was not clear how the SAF-A chromatin decondensation and RNA tethering activities were related. We now show that RNA:DNA tethering by SAF-A is performed by the monomeric form of the protein, and that mutations that trap SAF-A in an oligomeric state cannot tether RNA to DNA. Further, we show that recombinant, monomeric SAF-A is alone sufficient to tether RNA to DNA. Further, since all cellular SAF-A is complexed with RNA (Caudron-Herger et al., 2019) these results suggest that at least two distinct populations of SAF-A:RNA complexes are present in cells: a monomeric population that tethers RNA to DNA, and an oligomeric population that decondenses transcriptionally active chromatin. We show that failure to remove the RNA-tethering form of SAF-A from chromosomes during mitosis results in anaphase segregation defects. It is currently not clear how oligomerization of SAF-A is controlled during mitosis and this will require further investigation.

Our work has implications for how cells inherit transcriptional states from one cell cycle to the next. The theory of mitotic bookmarking holds that there are structures left in place that enable a chromosomal “memory” of the gene expression program prior to transcriptional downregulation in early mitosis. Then, as cells restart transcription in telophase, bookmarking structures – such as transcription factors or histone modifications – facilitate restoration of the gene expression program from the previous interphase, thereby reinforcing cell identity. Our images of RNA during a natural cell cycle revealed that cells evict nuclear and chromosome-bound RNAs concomitant with chromosome condensation and nuclear envelope breakdown, suggesting it is unlikely that there are RNA-based structural bookmarks associated with chromosomes that enable transcriptional memory. Instead, because we observed relocalization of both SAF-A and labeled RNA to chromosomes coincident with mitotic exit, it is possible there is a more nuanced role for SAF-A and RNA in transcriptional restart, such as enabling the decondensation of mitotic chromosomes as genes are reactivated. Further investigation will be needed to determine how the cell coordinates the return of SAF-A:RNA complexes to chromatin when mitosis is completed, and whether this facilitates chromosome remodeling during mitotic exit.

## Acknowledgements

We thank Brian Chadwick, Iain Cheeseman, Andrew Holland, and Paul Kaufman for sharing plasmids and cell lines. We are indebted to Jesse Cochrane and Radhika Subramanian for advice on performing the molecular modeling of SAF-A. We thank Iain Cheeseman, Paul Kaufman, Jeannie Lee, Carlos Perea-Resa, Barbara Panning, and Radhika Subramanian for insightful discussion and comments on the manuscript. This work was supported by a grant from the NIH to M.D.B. (GM122893).

## Author Contributions

Conceptualization, J.A.S. and M.D.B.; Methodology, J.A.S. and M.D.B.; Software, M.D.B.; Validation, J.A.S. and W.W.; Formal Analysis, J.A.S. and M.D.B.; Investigation, J.A.S., W.W., and M.D.B.; Data Curation, M.D.B.; Writing – Original Draft, J.A.S. and M.D.B.; Writing – Review and Editing, J.A.S. and M.D.B.; Visualization, J.A.S.; Supervision, J.A.S. and M.D.B.; Funding Acquisition, M.D.B.

## Declaration of Interests

The authors declare no competing interests.

---

**Supplemental Figure 1.**
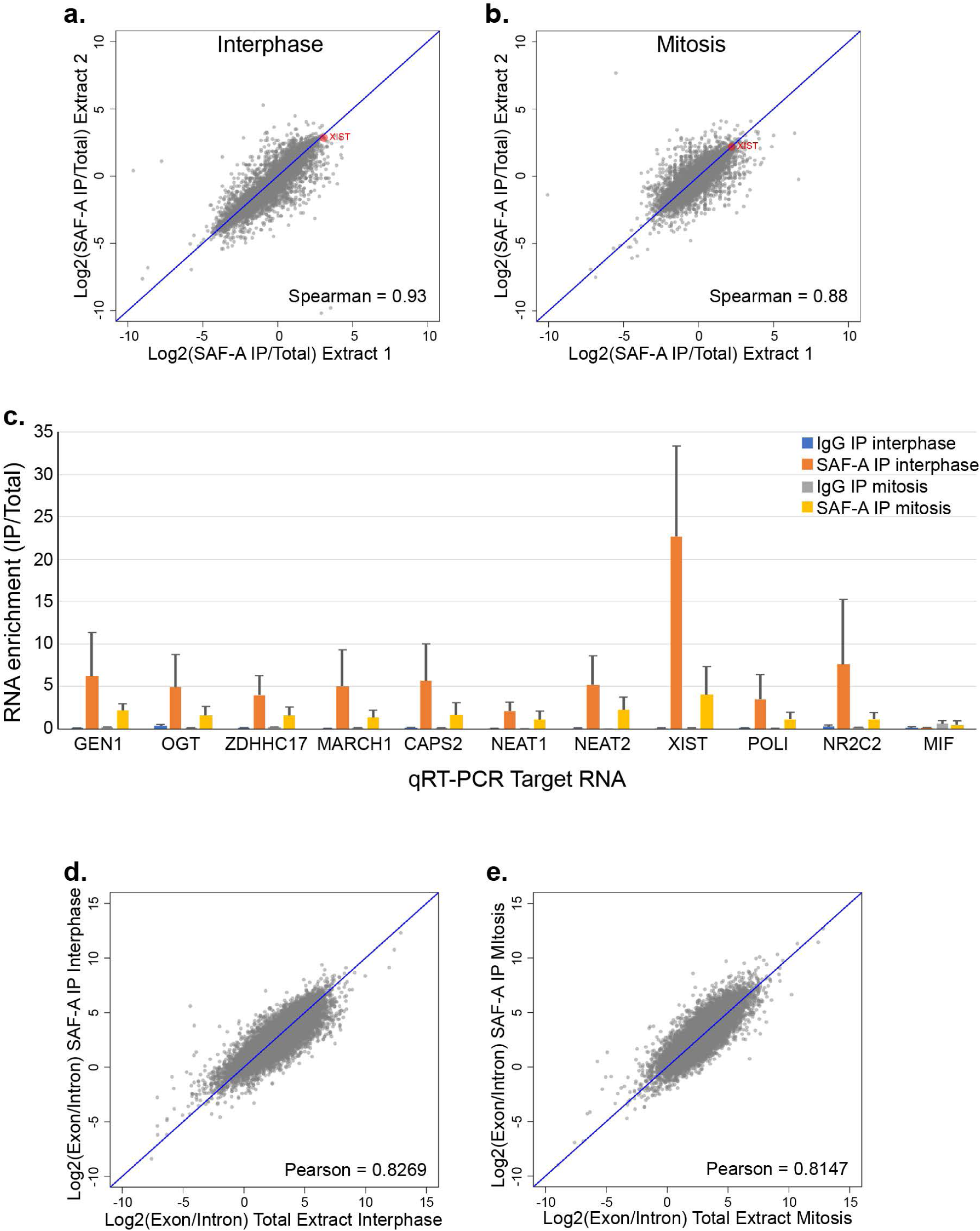
RIP-seq was performed to identify SAF-A-enriched RNAs in interphase and mitotic cell extracts; cDNA reads present in SAF-A IPs were aligned to the human genome. To determine the reproducibility of two biological RIP-seq experiments, we plotted RNA enrichment in SAF-A immunoprecipitations in the first experiment against RNA enrichment in the second experiment. **A-B.** Correlation plots for RIP-seq data from interphase extracts (**A**) and mitotic extracts (**B**). XIST RNA is highlighted in red. The Spearman correlation value is depicted on each plot. **C.** Quantitative RT-PCR of SAF-A-interacting RNAs identified by RIP-seq. All SAF-A-interacting RNAs tested showed significant enrichment in SAF-A IPs relative to their abundance in control immunoprecipitations. MIF was used as a negative control, since this RNA was not enriched in SAF-A IPs. SAF-A enrichment values represent the average obtained from two independent biological replicates; error bars represent the standard deviation. **D-E**. To determine whether SAF-A primarily interacted with mature or nascent RNA transcripts, we plotted the ratio of exon reads and intron reads for each transcript in the Total extract (x-axis) and SAF-A IPs (y-axis). Correlation plots are shown for RIP-seq data from interphase extracts (**D**) and mitotic extracts (**E**). Each point represents the average of two biological replicates. The Pearson correlation value is shown on each plot.

**Supplemental Figure 2.**
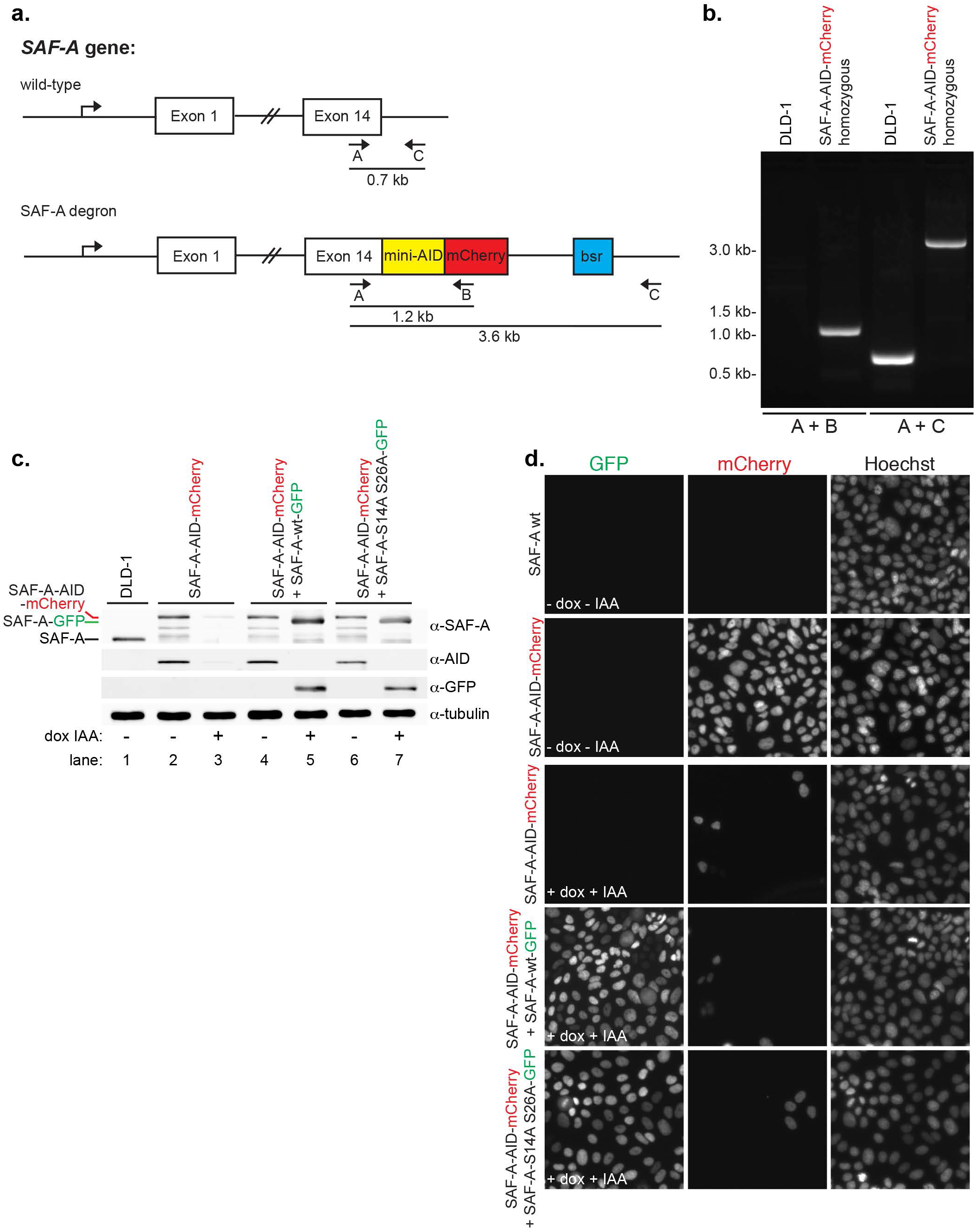
SAF-A depletion and reconstitution. **A.** The endogenous *SAF-A* locus was targeted for recombination in diploid DLD-1 cells, using CRISPR to introduce a C-terminal tag encoding both the minimal auxin-inducible degron sequence and mCherry. Clonal isolates were screened for homozygous SAF-A-AID-mCherry recombinants using the indicated combinations of primers A, B, and C. **B.** PCR analysis of genomic DNA from wild-type untransfected DLD-1 cells and the homozygous SAF-A-AID-mCherry clone used for depletion and reconstitution experiments **C.** Western blot analysis of SAF-A alleles in DLD-1 cells (lane 1), SAF-A-AID-mCherry cells (lanes 2-3), SAF-A-AID-mCherry + SAF-A-wt-GFP cells (lanes 4-5), and SAF-A-S14A S26A-GFP cells (lanes 6-7). SAF-A-AID-mCherry cells expressed the recombinant tagged protein at levels comparable to the wild-type untagged protein (lanes 1-2); addition of doxycycline and auxin to cell cultures resulted in extensive depletion of SAF-A-AID-mCherry (lane 3). The exclusive protein expression states represented in Figure 7A was confirmed for all tagged cell lines (lanes 2-7, compare – and + dox IAA) as depicted in the α-SAF-A, α-AID, and α-GFP blots. The relative mobility of all SAF-A protein species is indicated on the α-SAF-A blot. We note all tagged cell lines showed some degree of minor degradation bands. **D.** Live cell imaging of cell lines with and without doxycycline and auxin treatment. SAF-A-AID-mCherry was extensively depleted in all cell lines treated with doxycycline and auxin (+dox +IAA). In addition, induction of SAF-A-GFP allele expression was observed throughout cell populations containing the wild-type or phosphomutant SAF-A-GFP alleles.

**Supplemental Figure 3.**
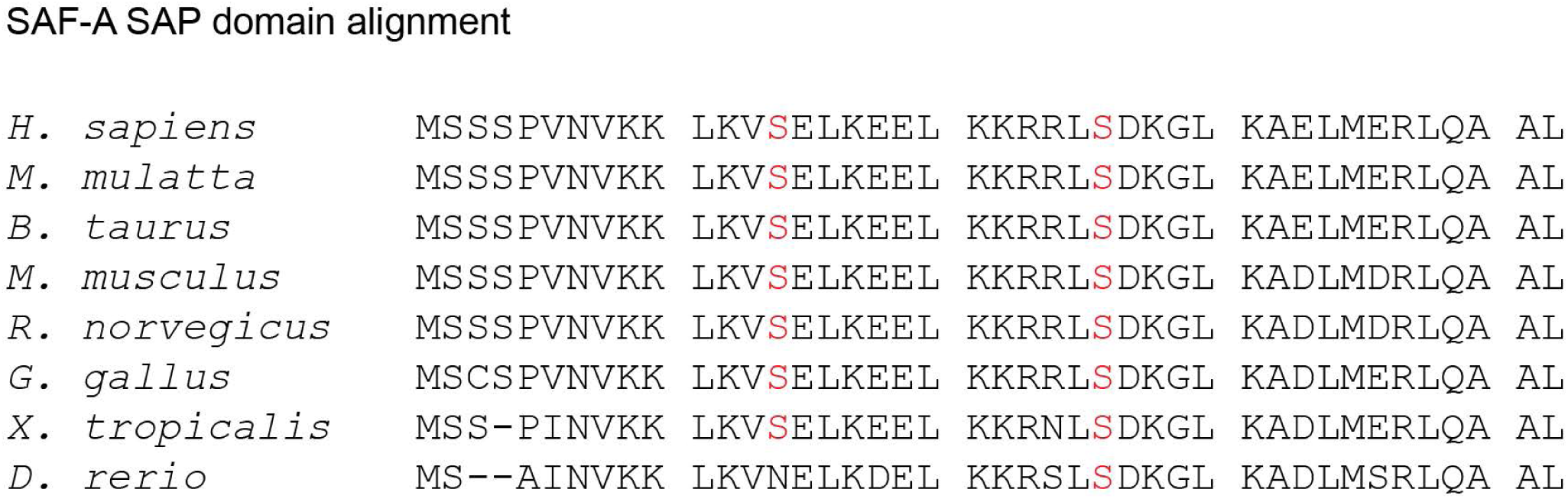
Alignment of the SAF-A SAP domain. SAF-A is conserved throughout vertebrates. Alignment of the SAF-A SAP domain indicates strong conservation of serines 14 and 26 throughout vertebrates, except in zebrafish SAF-A which retains only the second serine.

## Supplemental Tables

**Supplemental Table 1. SAF-A phosphopeptides identified in interphase and mitosis, from purification shown in Figure 4B.**

**Supplemental Table 2. RIP-seq data for SAF-A IPs in interphase and mitosis.**

## Methods

### Cell Culture

hTERT-RPE-1 cells were a gift from Brian Chadwick (Florida State University), and were cultured in DMEM-F12 supplemented with 10% FBS (Hyclone), Pen/Strep, 2 mM glutamine, and 7.5% sodium bicarbonate. 293T cells were a gift from Paul Kaufman (University of Massachusetts Medical School), and were cultured as recommended by the ATCC. DLD-1 cells were a gift from Andrew Holland. Cell lines were authenticated by microscopic observation of cell morphology, confirmation of the presence and number of Xi chromosomes, and were determined to be free of viral infection or mycoplasma contamination after testing by Charles River Laboratories.

### Cell synchronization and drug treatment

Cells were synchronized in interphase by incubating cells in 2 mM thymidine (Sigma, T-1895) for 24 hours. For synchronization in mitosis, cells were arrested first in a single thymidine block, washed twice in PBS and released into thymidine-free media for 6 hours, followed by an overnight incubation in 50 μM S-trityl-L-cysteine (Fluka, 93450). Cells were harvested the next morning by mitotic shake-off.

Barasertib (Mortlock et al., 2007) (Selleckchem, AZD1152-HQPA) and Aurora A inhibitor I (Aliagas-Martin et al., 2009) (Selleckchem, S1451) were each used at 1 μM. BI 2536 (Lenart et al., 2007) (Selleckchem, S1109) was used at 100 nM. An equivalent volume of DMSO alone was added to control cells. All drug stocks were prepared as 10 mM in DMSO and frozen until use. Cells were incubated in the presence of the drugs for 3-4 hours at 37°C prior to fixation.

### RNA labeling and detection

Total cellular RNA was labeled for 3 hours with EU (Thermo-Fisher, E10345) or BrU (Sigma, 850187). EU-RNA was detected using the Click-iT RNA Imaging Kit (Thermo Fisher, C10329 and C10330) according to the manufacturer’s instructions. When EU detection was combined with SAF-A immunofluorescence, we performed the SAF-A IF first, followed by a 10 minute fixation with 2% paraformaldehyde. Coverslips were then processed for EU-RNA detection as described above. SAF-A:RNA complexes were detected in BrU-labeled cell populations using the rabbit α-SAF-A and mouse α-BrU antibodies in conjunction with the Duolink Proximity Ligation Assay kit (Sigma, DUO92102).

### Immunofluorescence and FISH

Cells were washed briefly in PBS and fixed for 10 minutes in PBS containing 4% paraformaldehyde (Electron Microscopy Sciences) at room temperature. Cells were then gently extracted in PBS containing 0.5% Triton-X-100 for 20 minutes. In the course of this study we compared several fixation methods and found these conditions best for preserving mitotic chromosome structure, as well as SAF-A staining and RNA localization. In particular, we found extracting the cells in the presence of hypotonic cytoskeletal buffer caused extensive swelling of chromatin and suboptimal imaging of mitotic stages.

After fixation, cells were blocked briefly in PBS containing 1% RNase-free BSA (Ambion UltraPure BSA, AM2616) and 0.2% Tween. Cells were incubated with the primary antibody at 37°C, and washed three times in PBS + 0.2% Tween. Secondary antibody incubations and washes were performed similarly. Chromosomes were stained with 40 ng/mL DAPI. In experiments when FISH was performed after immunostaining, RNAsin (Promega) was added to the blocking buffer and antibody incubations at 400 U/mL, and DAPI staining was omitted. Prior to FISH, cells were fixed for 10 minutes in 2% paraformaldehyde to preserve antibody-antigen interactions throughout the FISH protocol.

For FISH, cells were fixed as described above and dehydrated sequentially in 70%, 85%, 95%, and 100% ethanol. The XIST RNA probe was directly labeled with Cy3-dUTP using Klenow (BioPrime DNA labeling system, Invitrogen) and a plasmid template with human XIST cDNA 1-16474 described previously (C. Xiao et al., 2007). Probes for the NEAT2 and OGT RNAs were labeled similarly, using a PCR product amplified from cDNA as the template. Cells were hybridized with the probe overnight in a humid chamber at 37°C in 50% formamide, 2X SSC, and 10% dextran sulfate. Coverslips were washed twice with 50% formamide/2X SSC, once with 2X SSC, and three times with 1X SSC. All washes were performed at 39°C. DAPI was included in the second 1X SSC wash at 40 ng/mL. Coverslips were mounted onto slides with Vectashield (Vector labs).

### Immunoprecipitation

After synchronization, cells were washed in PBS and incubated 30 minutes at 4°C in ice-cold lysis buffer (1 mL per 15 cm plate) containing 25 mM Tris pH 7.4, 150 mM KCl, 5 mM EDTA, 5 mM MgCl_2_, 1% NP-40, 0.5 mM DTT, and protease inhibitors (Roche). Cells were collected from the plate and passed several times through a syringe with a 22g needle. Lysates were centrifuged 30 minutes at 4°C (~22,000 × g) to remove insoluble material. Protein concentration of the extract was determined using a Bradford assay. Cell extracts were normalized to between 1-2 mg/mL, and incubated with 6 μg SAF-A antibody per mL extract for 1.5-2 hours at 4°C. Control immunoprecipitations included 6 μg rabbit or mouse IgG (Jackson ImmunoResearch) per mL extract, matched to the host species of SAF-A antibody used in experimental IPs. Immune complexes were collected 1.5-2 hours at 4°C, with pre-equilibrated Dynabeads (Thermo-Fisher; 4.5 μL Dynabeads per μg antibody) conjugated to either Protein A or Protein G, according to whether the antibody to SAF-A was generated in rabbit or mouse, respectively. The slurry of beads bound to immune complexes was washed three times in 1 mL lysis buffer prior to elution.

For immunoaffinity purification of SAF-A for mass spectrometry, we used a rabbit polyclonal antibody to SAF-A, and included protein phosphatase inhibitor cocktails I and III (Sigma-Aldrich) in cell extracts. Samples were submitted for analysis to the Taplin Biological Mass Spectrometry Facility (http://taplin.med.harvard.edu/). Phosphopeptides identified in this study for both interphase and mitotic SAF-A are detailed in Supplementary Table 1. The schematic of SAF-A phosphopeptides mapped relative to domain structure was assembled from this study,(Kettenbach et al., 2011; Olsen et al., 2010); www.phosphosite.org and references therein.

To investigate SAF-A-associated RNAs (RIP-seq), we performed immunoprecipitation as described above, except that a mouse monoclonal antibody to SAF-A was used. RNAsin (Promega) was added to the IP at 40 U/mL. Samples were eluted in Trizol (Thermo-Fisher) for RNA purification prior to library production. The co-precipitation of SAF-A with chromatin was also performed with the mouse monoclonal antibody to SAF-A. Samples were treated as described above, except that immune complexes were formed during an overnight incubation at 4°C.

### qRT-PCR

Superscript III reverse transcriptase (Invitrogen) was used to synthesize cDNA at 50°C according to the manufacturer’s instructions. For quantitative real-time PCR, we used 1/20^th^ volume of a cDNA reaction to program a PCR reaction using iQ SYBR Green Supermix (Bio-Rad). PCRs were amplified on a CFX96 Real-Time System (Bio-Rad) and analyzed with the CFX software package.

### Antibodies

Primary antibodies used in this study were: rabbit mouse α-SAF-A (Abcam, ab10297); rabbit α-SAF-A (Abcam, ab20666); rabbit α-histone H3 (Abcam, ab1791); mouse α-histone H3-S10P (Abcam, ab14955); mouse α-GFP (Abcam, ab1218); mouse α-Aurora-B (Abcam, ab3609) and a mouse antibody with reactivity to BrU ((Paulsen et al., 2014) BD Biosciences, 555627). SAF-A-AID-mCherry protein was detected using an alpaca α-RFP antibody conjugated to Atto-594. (Bulldog Bio, RBA594). Tet-inducible SAF-A-GFP alleles were detected using alpaca α-GFP (Bulldog Bio, GBA488).

### Transfections

Full-length, wild-type SAF-A was cloned into pEGFP-N1 (Clontech) and confirmed by DNA sequencing. The resulting plasmid encoding SAF-A (pMB918) was used as a template for site-directed mutagenesis to generate phosphomutant and phosphomimetic alleles of SAF-A. Plasmids used for transfections were: pMB918 (SAF-A^wt^-GFP), pMB935 (SAF-A^S14A^-GFP), pMB936 (SAF-A^S26A^-GFP), pMB1003 (SAF-A^S14A S26A^-GFP, pMB992 (SAF-A^S271A^-GFP), pMB1004 (SAF-A^S26A S271A^-GFP), pMB932 (SAF-A^S267A S271A^-GFP), pMB931 (SAF-A^S187A S188A T191A S192A^-GFP), pMB1013 SAF-A^S14D S26D^-GFP, and pMB1014 (SAF-A^S14E S26E^-GFP).

293T cells were plated onto coverslips the day before transfection at a density of 0.3 × 10^6^ cells per well. Cells were transfected for 48 hours with wild-type or mutant versions of SAF-A-GFP using 2.5 μg plasmid DNA and 2.0 μL Lipofectamine 3000 (Invitrogen). Three independent transfections were analyzed to assess the frequency of the phenotypes reported in Figures 4 and 5.

To deplete Aurora-B by RNA interference, RPE-1 cells were transfected using RNAiMAX (Invitrogen) and incubated for 48 hours prior to processing coverslips for FISH and immunostaining. Cells were transfected using a non-targeting siRNA (Dharmacon, D-001210-01-05) or a pool of siRNAs specific for Aurora-B (Dharmacon, M-013501-01-0005).

### Microscopy

Cells were visualized on an Olympus microscope (BX61) equipped with a DSU spinning disc confocal attachment. Images were captured as 3D optical stacks (0.2 μM per slice) using a charge-coupled device camera (ORCA-ER; C4742-80; Hamamatsu Photonics). A 100X oil-immersion lens (NA 1.4) was used for all images. Coverslips were mounted using Vectashield, and visualized under Olympus immersion oil. Images were captured at room temperature. Image stacks were processed and analyzed using the MetaMorph software package (Molecular Devices).

For image quantitation, we acquired image series for each experimental condition using equivalent exposure times. In each optical stack, we identified a single slice representing the midpoint of the chromosome rosette. We measured total EU-RNA or SAF-A fluorescence by subtracting background fluorescence, thresholding the image, marking the cell boundary, and then summing pixel intensity over the selected area. To measure chromatin overlap of EU-RNA and SAF-A, DAPI images were deconvolved and used to select the chromatin area by thresholding. We then summed pixel intensity of EU-RNA or SAF-A within the area overlapping chromatin. The chromosome-localized fluorescent signal was subtracted from the total intensity to obtain the cytoplasmic fluorescence. EU-RNA or SAF-A localization was then expressed as a ratio of chromosome overlap/cytoplasmic localization.

### Aurora-B Kinase Assay

Aurora-B:Incenp complex was expressed and purified from *E. Coli* as described (Jambhekar, Emerman, Schweidenback, & Blower, 2014). SAF-A was immunoaffinity purified from interphase cells, using the α-SAF-A rabbit polyclonal antibody. Kinase assays were performed as described (Bolton et al., 2002; Rosasco-Nitcher et al., 2008).

### Molecular modeling

The DNA binding SAP domain of SAF-A (residues 1-62) was submitted to the I-TASSER server (http://zhanglab.ccmb.med.umich.edu/I-TASSER/)(Roy, Kucukural, & Zhang, 2010; Yang et al., 2015) to identify structural homologs present in the protein data bank (http://www.rcsb.org). We chose the highest ranked of five models for further analysis. Phosphoserines were modeled onto the SAF-A SAP domain model using Vienna-PTM 2.0 (http://vienna-ptm.univie.ac.at(Margreitter, Petrov, & Zagrovic, 2013; Petrov, Margreitter, Grandits, Oostenbrink, & Zagrovic, 2013). Molecular models of wild-type and phospho-SAF-A were rendered in PyMOL (www.pymol.org). Electrostatic surface potential and surface charge calculations were rendered in PyMOL using the APBS plug-in (www.poissonboltzmann.org). *SAF-A SAP domain purification and DNA EMSA*

The wild-type SAP domain of SAF-A was amplified by PCR and cloned into a modified pET30a vector containing GFP using In-Fusion cloning (Clontech; pMB1018). SAP-S14D-S26D (pMB1028) was created using site-directed mutagenesis. Clones were verified by sequencing. Proteins were expressed in BL21 Rosetta 2 cells overnight at 18°C, and were lysed using a French press in PBS containing 300 mM NaCl and EDTA-free protease inhibitors. Lysates were cleared by centrifugation at 25,000 × g for 30 minutes at 4°C. Cleared lysates were bound to Ni-NTA resin (Qiagen) for 1 hour at 4°C. Beads were washed with ~100 column volumes of PBS containing 300 mM NaCl and 10 mM imidazole. Proteins were eluted using PBS containing 500 mM imidazole. Eluted proteins were immediately loaded onto a Superdex S200 column equilibrated with 50 mM Tris pH 7.5, 150 mM NaCl. Peak fractions were pooled, adjusted to 10% glycerol, and stored at −80°C.

The DNA template for EMSA was an AT-rich SAR sequence (Okubo et al., 2004). AAT TCA GAA AAT AAT AAA ATA AAA CTA GCT ATT TTA TAT TTT TTC) was ordered as DNA oligos from IDT. Oligos were annealed and end-labeled with *γ*-^32^P ATP using PNK. PNK labeling reactions were purified using a G50 spin column, then 1 μg of labeled DNA was gel purified from a 10% native PAGE gel. After elution, labeled DNA was ethanol precipitated and quantitated by Nanodrop. Purified DNA duplex was adjusted to 50 nM and stored at −20°C.

DNA EMSA reactions contained the indicated protein concentrations (Figure 5F), 5 nM annealed SAR oligos, in a buffer containing 50 mM Tris pH 7.5, 50 mM NaCl, 1 mM EDTA, and 4% glycerol. All components were mixed and incubated at room temperature for 30 minutes. Reactions were separated in a 10% native PAGE gel in 1X TBE run at 120 V for 1 hour. Gels were dried and exposed to a Phosphor screen overnight. Binding reactions were performed in duplicate. The fraction of the SAF-A SAP domain bound to the dsDNA SAR template was calculated using ImageJ, and binding curves were fit to the data using R.

### Expression and purification of SAF-A in Sf9 cells

Full-length SAF-A was cloned into pFB-HTA using InPhusion cloning (pMB1105). Sf9 cells were grown to a density of 1 × 10^6^ cells per ml and infected with a MOI ~5 from a P2 viral stock. Infected cells were grown for 48h after infection. Infected cells were collected by centrifugation at 1000 × g for 15 minutes. Cells were washed once with ice-cold PBS and collected by centrifugation as above. Cell pellets (~1 × 10^9^ cells) were resuspended in 25 mL PBS (400 mM NaCl final concentration), 1X protease inhibitors (Roche Complete Mini), benzonase (50U, Sigma), and 1% CHAPS. Resuspended cells were lysed by 10 passages of a tight-pestle Dounce homogenizer. Cells were centrifuged at 40,000 × g for 15 minutes at 4°C. A volume of 0.8 mL NiNTA (Qiagen) beads were added to the soluble lysate and incubated in batch at 4°C for 1 hour. Beads were collected and washed in column format with ~50 column volumes of PBS. Protein was eluted with 1 mL PBS + 50 mM imidazole. Eluted protein was diluted 2:3 with cold water. Eluted protein was applied directly to a HiTrap heparin column on a Bio-Rad FPLC. Bound protein was eluted with a NaCl gradient from 100 – 1000 mM NaCl. SAF-A eluted in a single peak from the heparin column. Peak fractions from the heparin column were loaded onto a Sephracyl S200 column run in 50 mM Tris pH 8.0, 100 mM NaCl, 0.5 mM EDTA. Peak fractions were collected and quantified by Bradford. The final yield was 1.5mg of purified SAF-A from 1.5 L of cultured Sf9 cells.

### DNA bead tethering assays

An 875 bp fragment of the MII SAR region (hg38 chr20:41104718-41105592) was amplified from human genomic DNA and cloned into pCR2.1 using TOPO cloning (pMB1156). This fragment was amplified from the plasmid using M13F-Biotin and M13R. Purified DNA was coupled to M280 streptavidin Dynabeads as described (Hannak & Heald, 2006) A 935 nt fragment from exon 1 of human XIST RNA (hg38 chrX:73845035-73845969) was prepared by *in vitro* transcription incorporating FITC-UTP. Transcribed RNA was purified by LiCl_2_ precipitation.

Tethering reactions were performed in 25 μL reaction volumes. We first prepared protein:RNA complexes by incubating purified SAF-A with XIST RNA. Reactions contained 100 nM XIST RNA, 2 μM SAF-A, 0.1 mg/mL yeast tRNA, 0.1% CHAPS in 50 mM Tris pH 8.0, 100 mM NaCl, 1 mM DTT. Reactions were incubated for 30 minutes at room temperature, then added to 15 μL of DNA (or empty streptavidin beads) and incubated for 30 minutes at room temperature with occasional flicking. Beads were washed 3 times, 1 mL per wash, in 50 mM Tris pH 8.0, 100 mM NaCl, 0.1% CHAPS, switching tubes once between washes 2 and 3. Protein:RNA:DNA complexes were eluted with 1X-SDS sample buffer by incubating at 80°C for 5 minutes. Half the eluted samples were run on agarose and acrylamide gels for RNA and protein analysis.

### SAF-A RIP-seq and bioinformatics

RNA associated with SAF-A was immunoprecipitated as described above, and purified in Trizol (Invitrogen). Any potential contaminating DNA was removed by DNase digestion (RQ-1, Promega); RNA was subsequently purified on a column (RNA Clean & Concentrator-25, Zymo Research). Ribosomal rRNA sequences were removed using a Ribo-Zero Gold rRNA removal kit (Illumina), the rRNA-free eluate was again column-purified (RNA Clean & Concentrator-5, Zymo Research). RNA libraries were generated using the NEBNext Ultra RNA Directional Library (New England Biolabs), PAGE purified, and sequenced using an Illumina HiSeq-2000 platform (nextgen.mgh.harvard.edu).

Reads from RNA-seq libraries were aligned to the human genome builds hg19 and hg38 using Tophat (Trapnell et al., 2012). Reads were counted against UCSC gene annotations (for hg38). To analyze differential gene expression we first filtered our RNA-seq data to eliminate any genes that had a FPKM of 0 in any of the 8 sequenced libraries. All plots were created using R. To identify coding and noncoding genes in the UCSC genome annotation we extracted a representative Refseq transcript for each gene annotation. We categorized genes as being a mRNA if the representative transcript had a name starting with “NM_” and noncoding RNA genes had the format “NR_.” Comparisons of noncoding RNA overrepresentation were calculated using a Fisher Exact Test in R. All sequences and summary tables generated by high-throughput sequencing have been deposited in GEO under accession number GSE87200. Fastq files and summary sequencing tables can be accessed during review using the private link: https://www.ncbi.nlm.nih.gov/geo/query/acc.cgi?token=ghuhkaawdtkxfwf&acc=GSE87200

To compare SAF-A RNA-seq data to nuclear retained RNAs(Djebali et al., 2012) we downloaded datasets of whole cell, nucleus, and cytoplasm RNAs from IMR-90 cells, a untransformed, female diploid primary cell line (SRR534291, SRR534292, SRR524299, SRR534300, SRR534301, SRR534302). Reads were aligned to hg38 using Tophat and counted against UCSC gene models using Cufflinks as described above. Readcounts were filtered for genes with a FPKM > 0 and combined with hnRNP-U IP-seq data.

### Statistics

Two or more independent biological replicates were performed for all experiments described in this study, with two exceptions. Identification of SAF-A phosphopeptides by mass spectrometry was performed once and pooled with published data as described above (Figure 4B-C). Molecular modeling of the SAF-A SAP domain was also performed once (Figure 5A-B). For statistical analysis of quantitative data, the central tendency is represented by the mean, and variation is represented by the standard deviation. *P*-values were calculated using a two-sided t-test assuming equal variance of the two samples; the number of individual data points (n) measured for each comparison is specified in the figure legends. Correlation statistics for RNA-seq experiments were calculated using Spearman’s correlation coefficient in R. Graphs were generated using Excel, Plotly, and R software.

### Data availability

All sequences and summary tables generated by high-throughput sequencing have been deposited in GEO under accession number GSE87200. Fastq files and summary sequencing tables can be accessed during review using the private link: https://www.ncbi.nlm.nih.gov/geo/query/acc.cgi?token=ghuhkaawdtkxfwf&acc=GSE87200

### Construction of cell lines expressing degron-tagged SAF-A

We targeted the endogenous SAF-A locus for C-terminal tagging using the CRISPR-Cas9 system as previously described (Natsume et al., 2016; Ran et al., 2013). In brief, we used the online CRISPR Design Tool (http://tools.genome-engineering.org) to identify a PAM sequence near the SAF-A stop codon and to design a corresponding guide RNA for cloning into the BbsI sites of pX330-U6-Chimeric_BB-CBh-hSpCas9 (Addgene #42230). The resulting construct, pMB1050, was used in conjunction with the targeting construct to modify the SAF-A locus.

To generate the SAF-A targeting construct, we fused 200 bp homology arms to a cassette encoding the minimal auxin-inducible degron sequence fused to the mCherry tag and the blasticidin-resistance gene. The 5’ homology arm corresponds to human chromosome 1, nucleotides 244854433-244854462. The 3’ homology arm corresponds to human chromosome 1, nucleotides 244854250-244854449. The tagging cassette was derived from plasmid pMK294 (mAID-mCherry2-Bsr; Addgene #72832). All elements of the targeting cassette were ligated together in the pBluescript KS+ vector (In-Fusion, Takara) to make pMB1116.

DLD-1 cells were cotransfected with pMB1050 and pMB1116, using the Lipofectamine 3000 transfection reagent (Thermo-Fisher). After 72 hours, cells were selected for six days with 5 μg/mL Blasticidin S (Thermo-Fisher). Individual clones were obtained by FACS sorting mCherry-positive cells at 1 cell/well of a 96-well plate. Cell lines were then screened by genomic DNA PCR, western blot analysis, and live cell imaging to identify positive clones expressing SAF-A-AID-mCherry (Figure S2).

SAF-A-AID-mCherry cells were transfected with a plasmid encoding codon-optimized, tet-inducible OsTIR1 (pMK243; Addgene #72835). Cells were selected with 0.5 mg/mL Geneticin (Thermo-Fisher); individual drug-resistant colonies were picked for expansion using cloning cylinders. Positive clones were first screened by RT-PCR to identify lines that express TIR1 after doxycycline treatment and were then tested for inducible degradation of SAF-A-AID-mCherry. We observed SAF-A degron cells required 24 hours of treatment with 1 μg/mL doxycycline (Sigma, D3447) and 500 μM auxin (3-indole-acetic acid, IAA, Sigma, I-5148) for optimal depletion of SAF-A-AID-mCherry.

### SAF-A-AID-mCherry depletion and reconstitution

GFP-tagged SAF-A alleles were cloned into the lentiviral expression vector pLVX-TetOne-puro (Takara) using In-Fusion cloning. Lentiviral plasmids were cotransfected with packaging plasmids pLP1, pLP2 and pVSV/G into 293T cells using Lipofectamine 2000 (Thermo-Fisher). Lentiviral-containing supernatant was collected at 72 hours and used to transduce SAF-A-AID-mCherry cells in the presence of RPMI media supplemented with 10% FBS and 6 μg/mL polybrene (Sigma, H-9268). After 48 hours, cells were passaged prior to FACS sorting of GFP-positive cells. Individual clones were then tested by western blot and live cell imaging to confirm SAF-A-GFP expression levels.

In experiments where SAF-A function was reconstituted with GFP-tagged inducible proteins, cells were first treated with 1 μg/mL doxycycline for 24 hours to induce TIR1 and SAF-A-GFP, followed by another 24 hour incubation in doxycycline and 500 μM IAA to degrade SAF-A-AID-mCherry.

